# Local ROS and inefficient autophagosome formation limit Optineurin-mediated mitophagy in neurons

**DOI:** 10.64898/2026.01.16.700014

**Authors:** Jordan R. Green, Anne E. Ojo, Molly K. Gooden, Lillian D. Papell, Titilola D. Kalejaiye, Chantell S. Evans

## Abstract

Mitophagy is a quality control mechanism that maintains neuronal health by removing damaged mitochondria. In neurons, mitophagy can be driven by Optineurin (OPTN), a selective autophagy receptor recruited in response to oxidative stress. However, mechanistic studies of mitophagy have relied mainly on non-neuronal cell lines and acute damage paradigms. It remains unknown whether these findings translate to neurons under physiological stress conditions. Here, we investigated the temporal dynamics of OPTN-dependent mitophagy under mild oxidative stress using live-cell imaging in primary rat hippocampal neurons. We found that autophagosomes failed to readily engulf OPTN-positive (OPTN+) mitochondria, revealing a previously unidentified rate-limiting step in neuronal OPTN-mediated mitophagy. Additionally, OPTN+ mitochondria colocalized with acidified lysosomes over timescales significantly longer than previously reported. These OPTN+ mitochondria displayed locally elevated reactive oxygen species (ROS). Scavenging mitochondrial ROS increased lysosomal colocalization with damaged mitochondria and accelerated mitophagy progression. Together, these findings indicate that redox signaling contributes to the slow timescale of OPTN-mediated mitophagy in neurons.

## INTRODUCTION

Mitochondria are dynamic organelles that undergo rearrangement and renewal to meet cellular needs. Quality control pathways preserve mitochondrial functional integrity through coordinated action (Jenkins et al., 2024; Misgeld and Schwarz, 2017). A key quality control mechanism is mitophagy, a selective form of autophagy. Mitophagy proceeds through coordinated steps that initially identify damaged mitochondria, followed by autophagosome sequestration and subsequent degradation within acidic lysosomes (Harper et al., 2018; Rodger et al., 2018; Wang and Klionsky, 2011; Youle and Narendra, 2011). To date, several parallel mitophagy pathways have been described (Evans and Holzbaur, 2020c; Villa et al., 2018), suggesting that cells may differentially regulate mitochondrial degradation in response to the type or severity of mitochondrial or cellular stress.

The best-characterized mitophagy pathway involves PTEN-Induced Putative Kinase 1 (PINK1) and Parkin. In this pathway, mitochondrial depolarization activates PINK1 and Parkin, leading to the formation of phospho-ubiquitin chains on outer mitochondrial membrane proteins (Kane et al., 2014; Kazlauskaite et al., 2014; Koyano et al., 2014; Matsuda et al., 2010; Narendra et al., 2008; Narendra et al., 2010; Ordureau et al., 2015; Vives-Bauza et al., 2010). These ubiquitin chains act as scaffolds for recruiting autophagy receptors such as Optineurin (OPTN). OPTN is highly expressed in the human brain and appears to be the predominant autophagy receptor driving neuronal mitophagy downstream of PINK1/Parkin (Evans and Holzbaur, 2020a; Lazarou et al., 2015). OPTN links ubiquitinated mitochondria to nascent autophagosome membranes through two key domains: the ubiquitin-binding in ABIN and NEMO (UBAN) and the microtubule-associated protein light chain 3B (LC3B) interaction region (LIR; (Gleason et al., 2011; Heo et al., 2015; Lazarou et al., 2015; Rogov et al., 2014; Stolz et al., 2014; Wild et al., 2011; Wong and Holzbaur, 2014; Zhu et al., 2007)). The engulfment of depolarized, ubiquitinated mitochondrial fragments into LC3B-studded autophagosomes facilitates subsequent fusion with lysosomes for eventual degradation (Youle and Narendra, 2011).

Most mitophagy studies have used immortalized, non-neuronal cell lines, such as HeLa, to examine the temporal dynamics of mitophagy. These studies relied on high concentrations of mitochondrial toxicants that generate reactive oxygen species (ROS). Elevated levels of ROS induce oxidative stress and mitochondrial damage (Guo et al., 2013; Kowaltowski and Vercesi, 1999). For example, the potent mitochondrial uncoupler carbonyl cyanide 3-chlorophenylhydrazone (CCCP) rapidly depolarizes mitochondria and increases ROS production (Fazli and Evans, 2023; Moore and Holzbaur, 2016). In cultured cells, overexpressed Parkin rapidly translocates to damaged mitochondria within minutes, and autophagosomes engulf OPTN-positive (OPTN+) mitochondria a few hours after damage (Wong and Holzbaur, 2014). These foundational studies in immortalized cell lines indicated that mitochondrial turnover is a rapid response that effectively removes dysfunctional mitochondria, thereby protecting the cell from damage. These experimental paradigms, however, may not fully reflect OPTN-mediated mitophagy dynamics under physiological mild stress in post-mitotic cells such as neurons.

Neurons are long-lived, highly polarized cells that rely heavily on mitochondria for functions such as calcium buffering and synaptic ATP production (Sheng and Cai, 2012). These demands place chronic stress on mitochondrial quality control mechanisms required for neuronal homeostasis and necessitate tight mitochondrial regulation. Thus, neurons may restrict mitophagy to preserve mitochondrial populations required to meet rapidly changing, localized demands (Li et al., 2021). This constraint likely slows mitochondrial sequestration and degradation in neurons, potentially shifting mitophagy kinetics relative to dividing cell lines.

Impaired mitophagy has been linked to neurodegeneration (Antico et al., 2025; Evans and Holzbaur, 2020c; Fang et al., 2019). Mutations in mitophagy-related genes, such as PINK1, Parkin, and OPTN, cause Parkinson’s disease (PD) and Amyotrophic lateral sclerosis (ALS), respectively (Kitada et al., 1998; Maruyama and Kawakami, 2013; Maruyama et al., 2010; Valente et al., 2004). Yet, most PD and ALS cases are sporadic (Byrne et al., 2011; Warner and Schapira, 2003), suggesting that, in addition to genetic mutations, intrinsic mechanisms contribute to neuronal vulnerability. The accumulation of damaged or dysfunctional mitochondria is a hallmark of neurodegenerative diseases. In neurons, inefficient or slow mitochondrial degradation over time may contribute to this accumulation. However, there is currently no consensus on neuronal mitophagy dynamics, and direct quantitative, comparative examination of individual steps of the pathway under physiologically relevant mitochondrial stress is lacking.

Here, we mechanistically characterized the temporal dynamics of OPTN-mediated mitophagy in primary hippocampal neurons using an antioxidant-deprivation paradigm as a mild physiological stressor. We combined this approach with live-cell imaging to temporally resolve individual steps within the pathway, from OPTN translocation to damaged mitochondria, through autophagosome engulfment, and degradation in acidic lysosomes. This approach enabled direct identification of a rate-limiting step in OPTN-mediated neuronal mitophagy under physiological stress. Our study shows that mitochondrial degradation in neurons takes days to complete and that this delay is due to inefficient autophagosome engulfment and local mitochondrial oxidative stress.

## RESULTS

### AOF treatment induces low levels of ROS and mitochondrial damage to promote OPTN-dependent mitophagy

We previously established an antioxidant-free (AOF) condition as a physiologically relevant mitochondrial-damaging paradigm to investigate OPTN-mediated mitophagy in neurons. A 6-hour AOF treatment induced low ROS levels, damaging a subset of mitochondria and restricting OPTN-dependent mitophagy to the soma (Evans and Holzbaur, 2020a). Based on these observations, we focused our current study on primary rat hippocampal neurons treated with control media that contain antioxidants, AOF, or 5 nM Antimycin A (AA, a potent mitochondrial electron transport chain inhibitor) for 6 hours to induce OPTN-dependent mitophagy. We visualized OPTN-mediated mitophagy within the neuronal soma using live-cell confocal microscopy and z-stack imaging (**Figure 1A**). We confirmed that AOF treatment induces ROS using CellROX, a fluorogenic probe. The CellROX fluorescence intensity in AOF-treated and AA-treated neurons was significantly increased compared to control, indicating elevated intracellular ROS (**Figure 1B-C**). Importantly, AOF produced substantially lower ROS than AA, supporting its use as a physiologically relevant, mild oxidative stress paradigm for characterizing the temporal dynamics of OPTN-mediated neuronal mitophagy.

**Figure 1:**
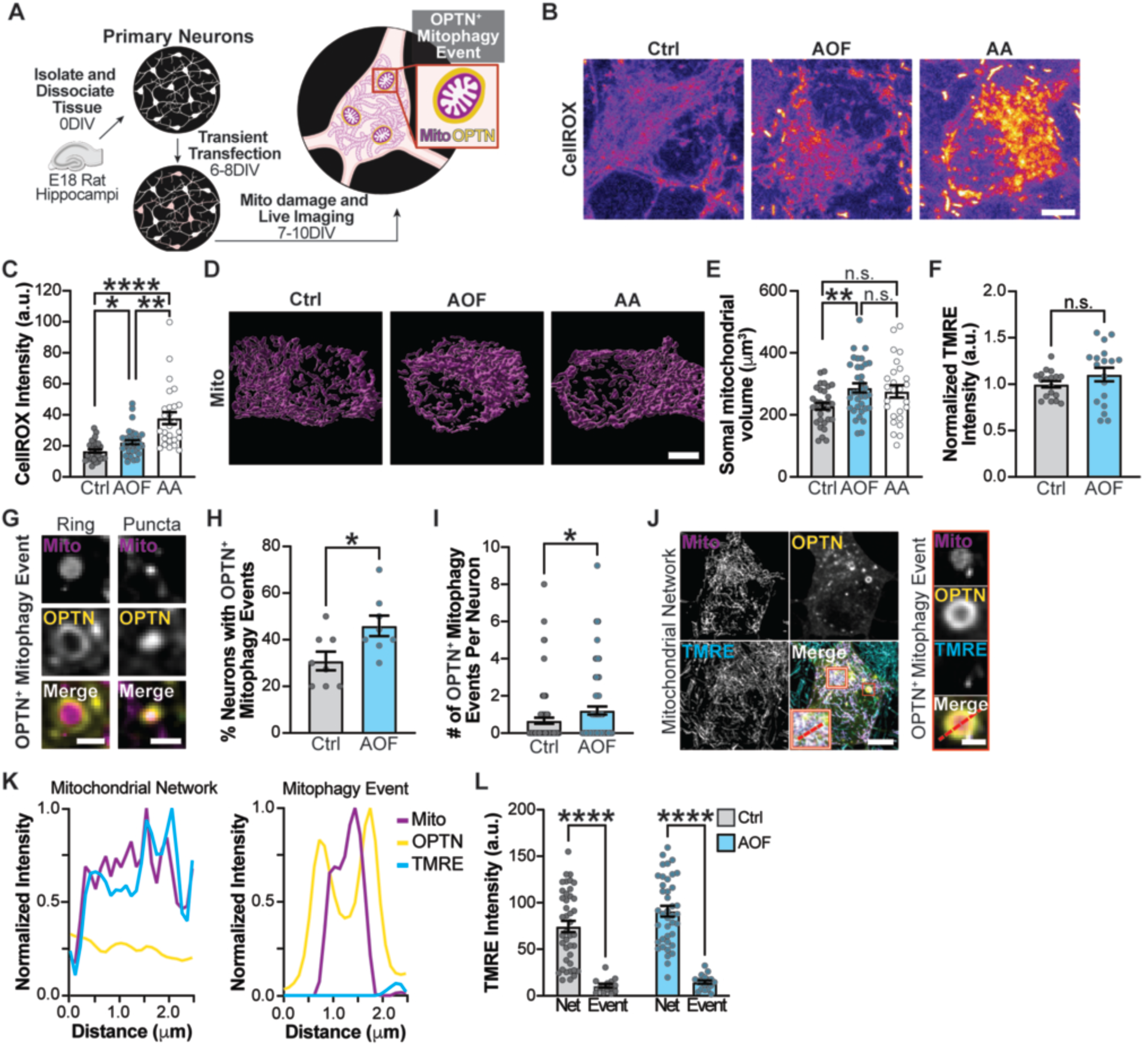
Antioxidant-free (AOF) medium induces mitochondrial dysfunction and OPTN-dependent mitophagy in primary hippocampal neurons. (**A**) Schematic of experimental paradigm using primary hippocampal neurons. Cultures were transfected prior to inducing mitochondrial damage, then the soma was live confocal imaged. (**B-C**) Representative images (B) and quantification (C) of intracellular ROS using CellROX Deep Red. Neurons were treated for 6 hours with either control, AOF, or 5 nM AA. Scale bar, 5 µm. Mean ± SEM; *n* = 26-35 neurons from 3 biological replicates; 8 DIV. *, *p* < 0.05; **, *p* < 0.01; ****, *p* < 0.0001 by Kruskal-Wallis with Dunn’s multiple comparisons test. (**D-E**) Representative images (D) and quantification (E) of the somal mitochondrial volume. Scale bar, 5 µm. Mean ± SEM; *n* = 26-35 neurons from 3 biological replicates; 8 DIV. n.s., not significant; **, *p* < 0.01 by Brown-Forsythe and Welch’s ANOVA with Dunnett T3 multiple comparisons. (**F**) Quantification of the TMRE fluorescence intensity. Mean ± SEM; *n* = 6 technical replicates per condition from 3 biological replicates; 8 DIV. n.s., not significant by Welch’s t-test. (**G**) Representative images of an OPTN+ mitophagy event; OPTN rings and puncta colocalized with fragmented mitochondria. Scale bar, 1 µm. (**H**) Quantification of the percent of neurons with OPTN+ mitophagy events. Mean ± SEM; *n* = 54-92 events from 8 plates per condition, which includes 76-81 neurons across 4 biological replicates; 8 DIV. *, *p* < 0.05 by unpaired t-test. (**I**) Quantification of the number of OPTN+ mitophagy events per neuron. Mean ± SEM; *n* = 54-92 events from 76-81 neurons across 4 biological replicates; 8 DIV. *, *p* < 0.05 by unpaired Mann-Whitney test. (**J**) Representative images of TMRE fluorescence signal in the mitochondrial network (left) and at an OPTN+ mitophagy event (right). The dashed red lines delineate the regions of interest used for the line scan intensity profiles in panel J of the mitochondrial network (inset, red and white box) and OPTN+ mitophagy event (red box). Scale bar: 5 µm (left); 1 µm (right). (**K**) Intensity profiles from the representative images in panel J, where distance refers to position along the line scan. At the OPTN+ mitophagy event, there is a specific loss of the TMRE fluorescence. (**L**) Quantification of TMRE fluorescence intensity in the mitochondrial network and at OPTN+ mitophagy events. Mean ± SEM; *n* = 14-17 events from 40-41 neurons across 4 biological replicates; 7 DIV. ****, *p* < 0.0001 by Two-Way ANOVA with Šidák’s multiple comparisons test.

We next assessed the effects of AOF treatment on mitochondrial structure and function, as well as on OPTN-mediated mitophagy induction. Neurons were transfected with a mitochondrially targeted BFP (Mito-TagBFP) to examine mitochondrial morphology and were co-labeled with the tetramethylrhodamine ethyl ester (TMRE) dye to measure mitochondrial membrane potential. While there was a small increase in mitochondrial volume following AOF treatment, there were no differences in overall TMRE fluorescence intensity (**Figure 1D-F**). Thus, AOF treatment induces subtle yet significant increases in ROS without globally affecting mitochondrial network function.

At mitochondrial fragments, we previously visualized OPTN structures that appeared as rings or puncta after 6 hours of AOF treatment, but focused our analysis only on OPTN rings (Evans and Holzbaur, 2020a). Here, we broadened our analysis to include rings and puncta, which we called OPTN-mediated mitophagy events (**Figure 1G**). We also observed mitophagy events under control conditions, attributable to constitutive mitophagy (McWilliams et al., 2016). AOF treatment significantly increased both the percentage of neurons with OPTN-mediated mitophagy and the number of mitophagy events per neuron compared to the control (**Figure 1H-I**). Therefore, OPTN-mediated neuronal mitophagy occurs under control conditions, and AOF induces mitochondrial ROS and increases the frequency of OPTN-mediated mitophagy, providing a useful tool for studying this pathway under physiological stress.

We confirmed that OPTN translocation was specific to damaged mitochondria with decreased membrane potential. Neurons were transfected with a mitochondria-targeted SNAP-tag (Mito-SNAP) and a Halo-tagged OPTN (Halo-OPTN) before loading the mitochondrial network with TMRE. We quantified TMRE fluorescence within the mitochondrial network and compared it with the TMRE signal of mitochondria at OPTN+ mitophagy events (**Figure 1J-L**). The TMRE signal was significantly higher in the intact mitochondrial network than at OPTN+ mitophagy events. These findings show that OPTN is specifically recruited to depolarized, damaged mitochondria, thereby serving as a visual marker of OPTN-mediated neuronal mitophagy.

### Autophagosomes fail to form in OPTN-mediated neuronal mitophagy

Having established that the AOF triggers OPTN-mediated mitophagy, we next sought to define the mechanisms underlying the temporal dynamics of this pathway in neurons. Mitophagy takes only a few hours to complete in non-neuronal, immortalized cell lines (Evans and Holzbaur, 2020a; Moore and Holzbaur, 2016). However, the time course and rate-limiting steps in neurons remain unclear. To resolve the individual stages of OPTN-mediated mitophagy, we used OPTN, LC3B, and Cathepsin D (CatD) or LysoTracker (LysoT) to reliably visualize autophagy receptor translocation, autophagosome formation, and lysosome degradation, respectively (**Figure 2A-B**). Based on our prior observation that Parkin and OPTN translocate to damaged mitochondria within 6 hours of AOF treatment (Evans and Holzbaur, 2020a), we focused our analysis on this time point to identify steps that might restrict mitochondrial turnover in neurons.

**Figure 2:**
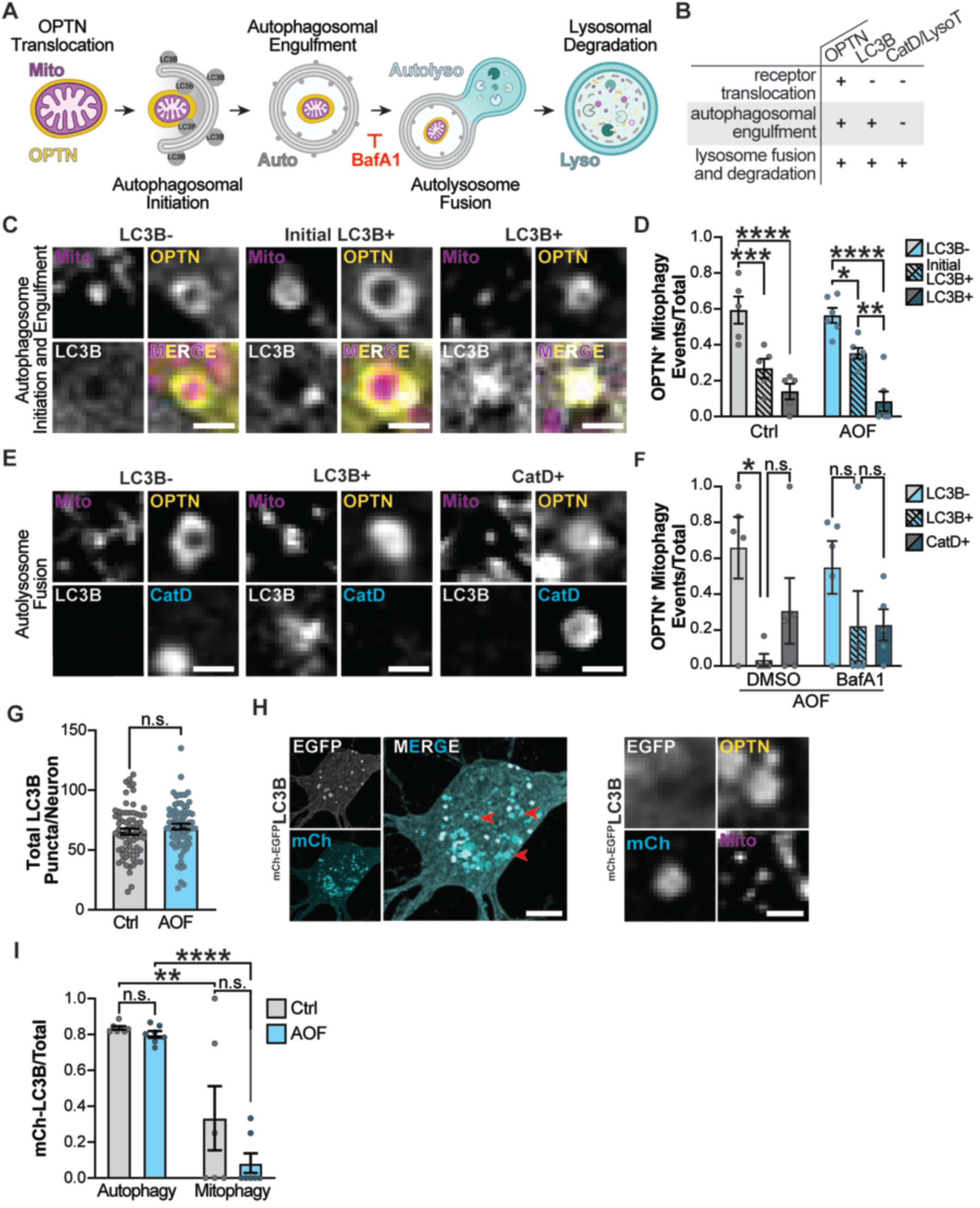
Inefficient autophagosome engulfment is rate-limiting in OPTN-mediated neuronal mitophagy. (**A**) Schematic of OPTN-dependent mitophagy. Briefly, autophagosome initiation and elongation engulf the OPTN+ mitochondrion. Once fully formed, the autophagosome fuses with the lysosome, generating an autolysosome; this step is inhibited by BafA1. Finally, lysosomal enzymes degrade the damaged mitochondrion. (**B**) Table illustrating the vesicular compartment that sequesters OPTN+ mitochondria and the markers used to identify each compartment. (**C-D**) Representative images (C) and quantification (D) of the fraction of OPTN+ mitophagy events that are LC3B-, Initial LC3B+, and LC3B+ following a 6-hour treatment with control or AOF. Scale bar, 1 µm. Mean ± SEM; *n* = 47-114 events from 5-6 plates per condition, which includes 45-52 neurons across 3 biological replicates; 8 DIV. *, *p* < 0.05; **, *p* < 0.01; ***, *p* < 0.001; ****, *p* < 0.0001 by Two-Way ANOVA with Tukey’s multiple comparisons test. (**E-F**) Representative images (E) and quantification (F) of the fraction of OPTN+ mitophagy events that are LC3B-, LC3B+, or CatD+ following a 6-hour AOF treatment plus DMSO or BafA1. Here, LC3B-encompasses both non-engulfed and initially engulfed LC3B events; CatD+ includes all OPTN-mediated events with a lysosome present, including the rare LC3B+/CatD+ events. Scale bar, 1 µm. Mean ± SEM; *n* = 24-27 events from 5 plates per condition, which includes 50-52 neurons across 5 biological replicates; 8-9 DIV. n.s., not significant; *, *p* < 0.05 by Two-Way ANOVA with Tukey’s multiple comparisons test. (**G**) Quantification of the total number of LC3B+ puncta per neuron. Mean ± SEM; *n* = 71 neurons from 7 biological replicates; 8-9 DIV. n.s., not significant by unpaired t-test. (**H**) Left, representative image of the dual-labeled mCherry-EGFP-LC3B autophagosome population. Most have undergone lysosomal fusion and acidification, as shown by the presence of cyan-only (mCherry+) puncta (red arrows). Scale bar, 5 µm. Right, representative image of an OPTN+ mitophagy event that colocalizes with an acidified autophagosome (mCherry+LC3B puncta). Scale bar, 1 µm. (**I**) Quantification of the fraction of mCherry+ LC3B puncta within autophagosomes or OPTN+ mitophagy events. Mean ± SEM; *n* = 17-29 events from 6-7 plates per condition, which includes 69-71 neurons across 7 biological replicates; 8-9 DIV. n.s., not significant; **, *p* < 0.01; ****, *p* < 0.0001 by Two-Way ANOVA with Tukey’s multiple comparisons test.

To evaluate autophagosome engulfment, neurons were transfected with mitochondrial-targeted DsRed2 (Mito-DsRed2), Halo-OPTN, and an EGFP-tagged LC3B (EGFP-LC3B). We first identified fully formed OPTN rings and puncta containing fragmented mitochondria. Then, we classified three types of OPTN+ mitophagy events: 1) those that were LC3B-negative (LC3B-); 2) those in which LC3B was targeted and bound to OPTN that appeared as a small punctum (Initial LC3B+); or 3) events where the LC3B punctum expanded to form a full autophagosome that engulfed the damaged mitochondria (LC3B+; **Figure 2C**). We quantified the proportion of events within each category relative to the total number of OPTN+ mitophagy events. In control and AOF conditions, ∼0.6 (60%) of OPTN+ mitophagy events remained LC3B-, suggesting that autophagosome initiation had not yet occurred. Moreover, only ∼0.15 (15%) of events were fully engulfed within an autophagosome (**Figure 2D**). The predominance of OPTN+ mitophagy events lacking LC3B at this time point indicates that autophagosome initiation and engulfment are markedly slow, suggesting a mechanistic delay in the pathway in hippocampal neurons.

Since EGFP-LC3B fluorescence is quenched in acidified compartments, it is possible that a subset of OPTN+ mitophagy events within lysosomes was not detected. To account for this, we used Bafilomycin A1 (BafA1), a known inhibitor of autolysosome fusion and acidification (**Figure 2A**). Neurons were transfected with Mito-SNAP, Halo-OPTN, EGFP-LC3B, and an RFP-tagged CatD (CatD-RFP), then treated with AOF and either BafA1 or a DMSO control for 6 hours. As expected, BafA1 treatment inhibited autolysosome fusion, significantly increasing the number of LC3B puncta while decreasing CatD puncta (**Figure S1A-B**). We identified OPTN+ mitophagy events that colocalized with LC3B or CatD (**Figure 2E**). Since the EGFP-LC3B fluorescence signal is quenched in acidic compartments, we used the loss of the EGFP signal as a proxy for mitophagy events that progressed into CatD-positive (CatD+) acidified lysosomes. We quantified the fractions of OPTN+ mitophagy events that were not engulfed (LC3B-), fully engulfed within an autophagosome (LC3B+), or contained within an acidified lysosome (CatD+). Consistent with previous data, most OPTN+ mitophagy events were LC3B-in both DMSO and BafA1 conditions (**Figure 2F**). These results indicate that 6 hours after treatment, most OPTN+ mitophagy events have not yet been engulfed by an autophagosome and thus lie upstream of BafA1-affected autolysosome fusion and acidification. We rarely observed OPTN+ mitophagy events within an autolysosome that lacked acidification (LC3B+/CatD+; 0.06 [6%]). While AOF increased OPTN-dependent mitophagy in percentage of neurons and number of events (**Figure 1H-I**), it did not alter the temporal dynamics compared with the control.

### Autophagosome initiation and formation are selectively impaired at OPTN+ mitophagy events

Since our data suggest that inefficient autophagosome initiation and engulfment limit OPTN-dependent mitophagy progression in neurons, we quantified autophagosomes within the soma. Neurons were transfected with Mito-SNAP, Halo-OPTN, and a dual-labeled mCherry-EGFP-tagged LC3B (mCherry-EGFP-LC3B). The EGFP fluorescence quenches in acidic environments, so the dual-labeled LC3B reporter enables simultaneous monitoring of autophagosome engulfment and acidification. We observed no significant difference in the total number of neuronal autophagosomes (LC3B puncta) or the fraction of acidified autophagosomes (mCherry-only LC3B puncta) between the 6-hour control and AOF conditions (**Figure 2G-I**). These data indicate that neuronal autophagosome abundance or their ability to fuse with acidified lysosomes is not altered.

We next compared the acidification of autophagosomes that colocalized with OPTN+ mitochondria (i.e., mitophagy) to those that did not (i.e., autophagy). We observed a significant decrease in the fraction of mCherry+ LC3B puncta that colocalized with OPTN+ mitochondria compared to those that did not (**Figure 2H-I**). Importantly, there was no difference in the fraction of acidified OPTN+ mitophagy events between the control and AOF conditions (**Figure 2I**; mitophagy, *p* = 0.1983). These findings indicate that autophagosomes involved in bulk autophagy form and acidify readily in neurons. In contrast, autophagosomes stall specifically at OPTN-mediated mitophagy events, suggesting that local features of damaged mitochondria hinder efficient autophagosome initiation and engulfment.

### Autophagosome initiation and formation limit the lysosomal degradation of OPTN+ mitophagy events

Our findings demonstrate that delayed autophagosome initiation and engulfment are rate-limiting in OPTN-mediated mitophagy in neurons. To confirm that the mitophagy progression stalls upstream of lysosomes, we quantified the fraction of OPTN+ mitophagy events that colocalized with LysoT, a dye that labels acidic lysosomes. Neurons were transfected with Mito-DsRed2 and Halo-OPTN and labeled with LysoT at the end of the 6-hour AOF treatment. We visualized OPTN+ mitochondria and found events that were either outside of a lysosome (LysoT-) or within (LysoT+; **Figure 3A**). Consistent with previous data, the vast majority of OPTN+ mitophagy events were LysoT- (**Figure 3B**). These data provide additional evidence that, in neurons, OPTN-mediated mitophagy rarely progresses to acidified lysosomes within 6 hours.

**Figure 3:**
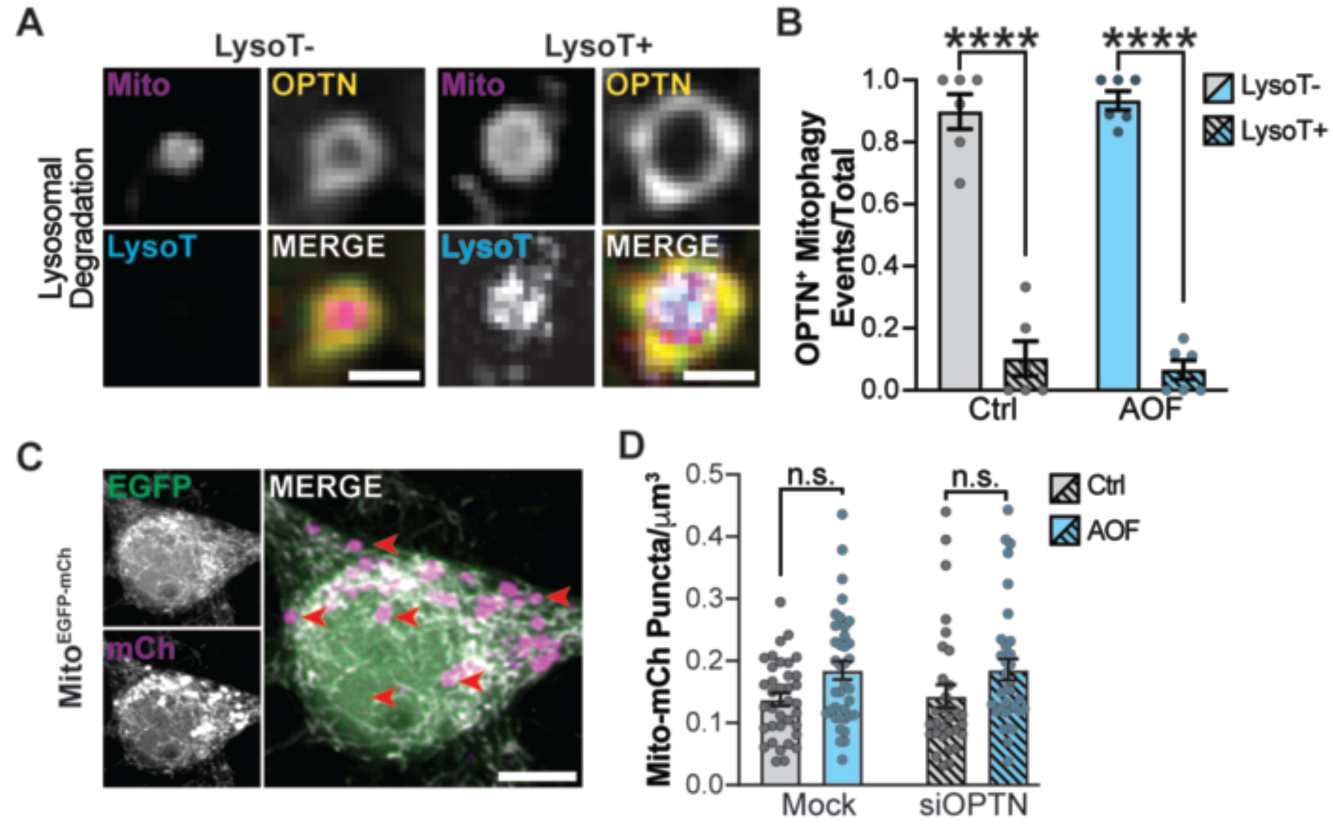
Impaired autophagosome engulfment prevents downstream lysosomal degradation of OPTN+ mitophagy events. (**A-B**) Representative images (A) and quantification (B) of the fraction of OPTN+ mitophagy events that are LysoT- and LysoT+ following 6 hours of control or AOF. Scale bar, 1 µm. Mean ± SEM; *n* = 32-54 events from 6 plates per condition, which includes 54-58 neurons across 3 biological replicates; 9 DIV. ****, *p* < 0.0001 by Two-Way ANOVA with Šidák’s multiple comparisons test. (**C-D**) Representative images (C) and quantification (D) of the number of mCherry+ mitophagy puncta per mitochondrial somal volume. Red arrows highlight acidified mitophagy events. Mean ± SEM; *n* = 31-37 neurons across 34 biological replicates; 9 DIV. n.s., not significant by Two-Way ANOVA with Tukey’s multiple comparisons test.

Next, we determined whether this delay extends broadly to all mitophagy pathways. We transfected neurons with the dual-labeled Mito-EGFP-mCherry mitophagy reporter. Mitophagy events are visualized as mCherry-only mitochondrial puncta (EGFP-/mCherry+) since EGFP fluorescence is quenched in the acidified lysosomes ((McWilliams et al., 2016); **Figure 3C**). Following a 6-hour treatment, mCherry+ mitochondrial puncta were quantified and normalized to mitochondrial volume. We observed a trending increase in the number of mCherry+ mitochondrial puncta following AOF treatment compared to control (**Figure 3D**). Since this is a general mitophagy reporter, the quantified events are likely a summation of multiple parallel mitophagy pathways. We visualized a small portion of OPTN+ mitophagy events colocalized with LysoTracker (**Figure 3A-B**), suggesting that OPTN-mediated mitophagy contributes only a small fraction of the events captured by the general mitophagy reporter. To test this idea, we used an siRNA to transiently knock down OPTN (**Figure S1C-D**) and quantified mCherry+ puncta. We found no differences between mock and siOPTN for either control or AOF (**Figure 3D**). These findings support our model and suggest that OPTN-mediated mitophagy has not progressed to lysosomal degradation within 6 hours. However, reduced expression of individual autophagy receptors correlates with increased abundance of parallel receptors, while inhibition of a single mitophagy pathway can alter the activity of parallel pathways (Lazarou et al., 2015; Puri et al., 2019; Yun et al., 2014). Thus, these mechanisms may contribute to our results.

Finally, we tested whether the delayed OPTN+ mitophagy we observed in control and AOF hippocampal neurons was neuron-specific. We previously found that acute treatment with Antimycin A and Oligomycin A (AA/OA) efficiently drives mitophagy progression, with OPTN+ mitochondria colocalizing with LysoT within a few hours (Evans and Holzbaur, 2020a). We therefore asked whether a milder stress would reveal similar OPTN-mediated mitochondrial dynamics in non-neuronal cells. Since our AOF paradigm cannot be directly translated to HeLa cells, we used a low dose of AA/OA (0.5 μM) to induce mild stress equivalent to AOF and compared it to severe stress (4 μM AA/OA) using CellROX (**Figure S1E-F**). HeLa cells were transfected with Mito-DsRed2 and Halo-OPTN and labeled with LysoT at the end of the 2-hour treatment. Most OPTN+ mitophagy events (∼0.6 or 60%) colocalized with LysoT under both mild and severe stress conditions (**Figure S1G-H**). Therefore, even under mild stress, OPTN+ mitophagy events turn over rapidly in non-neuronal cells, whereas this process is delayed in neurons.

### “Aged” mitochondria remain in hippocampal neurons for days after OPTN-mediated mitophagy induction

We next defined the full timescale of OPTN-mediated mitophagy in neurons. Previous studies have reported strikingly different mitophagy timescales in hippocampal neurons, ranging from minutes to days (Ashrafi et al., 2014; Lin et al., 2017). However, harsher reagents and varying doses were used to induce mitochondrial damage, potentially altering the kinetics of neuronal mitophagy. Therefore, we used our AOF paradigm to probe the temporal dynamics of mitophagy under physiological oxidative stress.

First, we examined the degradation of mitophagy-associated proteins Parkin, OPTN, LC3B-I, LC3B-II, and the mitochondrial marker Cytochrome C (CytC) via Western blot. LC3B-II is the lipidated form of LC3B-I and is associated with autophagosome formation. Neurons were treated with control, AOF, or AA and then incubated with cycloheximide (CHX), a protein synthesis inhibitor. Thus, any potential changes in protein abundance would reflect degradation. The expression of all mitophagy-associated proteins remained relatively stable across treatments and time points (**Figure 4A-B**). Since AOF treatment significantly increased the percentage of neurons with OPTN+ mitophagy and the number of OPTN+ mitophagy events compared to control (**Figure 1H-I**), these findings suggest that mitochondrial degradation via mitophagy is incomplete within 24 hours of mild oxidative stress. We note that changes in protein abundance due to degradation may be difficult to detect using bulk immunoblotting, as the fraction of total protein turnover attributable to mitophagy is relatively small (**Figure 1I**).

**Figure 4:**
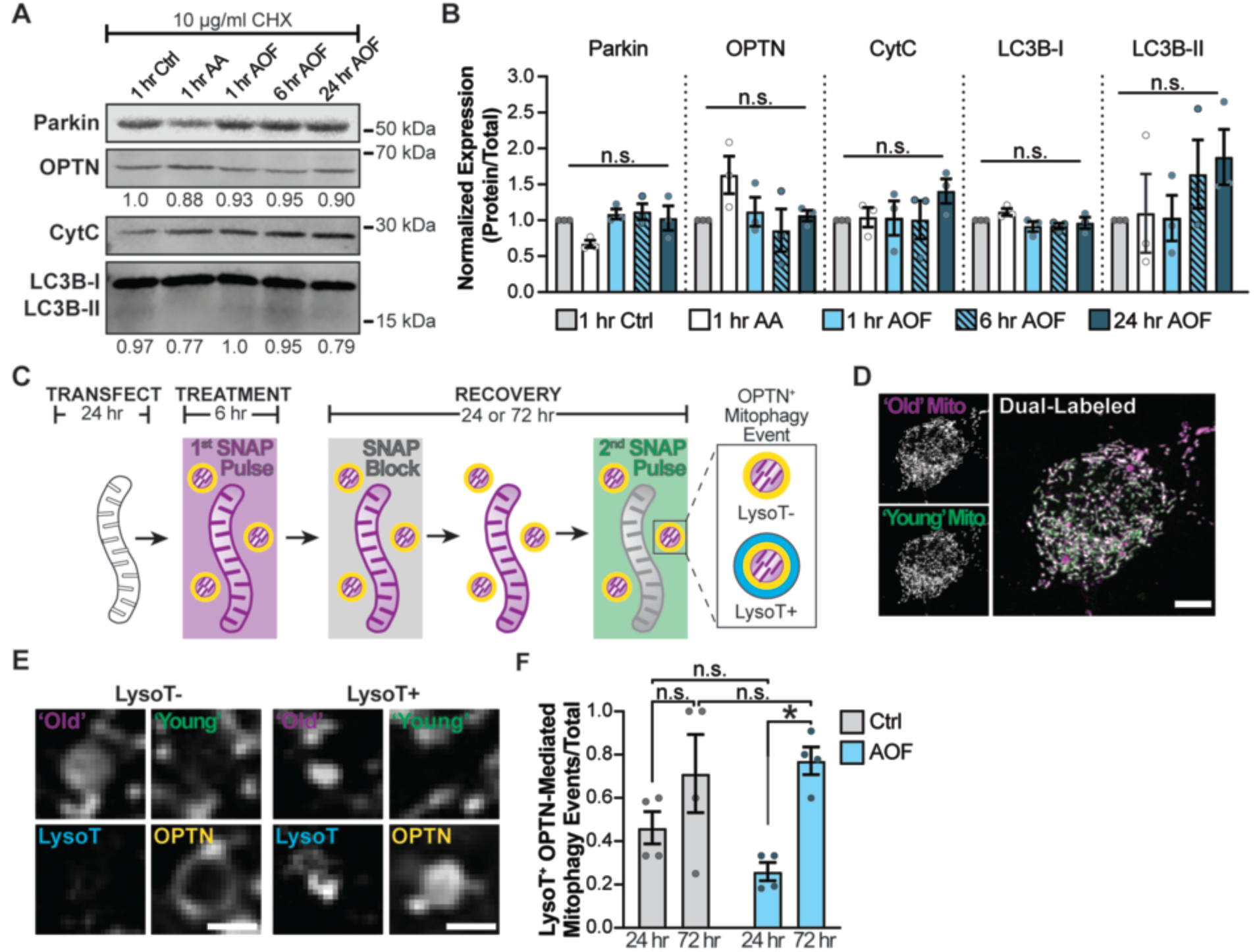
OPTN-mediated mitophagy takes days to fully complete in primary hippocampal neurons. (**A-B**) Representative fluorescent western blots (A) and quantification (B) of mitophagy-associated proteins following mitochondrial damage. Neurons were treated with Ctrl, AA (100 nM), or AOF for varying time points, with 10 µg/mL cycloheximide (CHX) added to inhibit protein synthesis. Normalization factors are shown under the representative images. Data shown as fold change over control for the protein of interest divided by the total protein stain. Mean ± SEM; *n* = 3 biological replicates; 8 DIV. n.s., not significant by One-Way ANOVA with Dunnett’s multiple comparisons test. (**C**) Schematic of the SNAP pulse-chase assay. Briefly, transiently transfected neurons were treated in parallel with AOF to induce OPTN-mediated mitophagy and labeled with the first SNAP pulse (magenta) for 6 hours. After the treatment, neurons were labeled with SNAP Block, followed by a 24- or 72-hour recovery period. In the final hour of recovery, neurons were labeled with the second SNAP pulse (green) and LysoTracker to visualize acidified lysosomes. (**D**) Representative images of spectrally distinct ‘old’ (magenta) and ‘young’ (green) mitochondrial populations, where most of the network is dual-labeled. Scale bar, 5 µm. (**E-F**) Representative images (E) and quantification (F) of the fraction of “old” OPTN+ mitophagy events that are LysoT+. Scale bar, 1 µm. Mean ± SEM; *n* = 24-60 events from 4 plates per condition, which includes 48-49 neurons across 3 biological replicates; 8-10 DIV. n.s., not significant; *, *p* < 0.05 by Two-Way ANOVA with Tukey’s multiple comparisons test.

For this reason, we directly visualized long-term mitochondrial turnover using a novel SNAP pulse-chase assay, enabling longitudinal tracking of individual mitochondria with resolution to capture discrete OPTN-mediated mitophagy events over days. This paradigm used sequential labeling with spectrally distinct SNAP ligands at defined time points to distinguish pre-existing ‘Old’ mitochondrial populations from newly synthesized ‘Young’ populations (Evans and Holzbaur, 2020a; Schneider and Evans, 2022). We first confirmed that the SNAP ligands bind the SNAP-tag fusion protein with equal propensity when added simultaneously (**Figure S2A**). Neurons were transfected with Mito-SNAP and Halo-OPTN and labeled with SNAP 430 and SNAP JF549 ligands. Under these conditions, the mitochondrial network and 96.5% of the OPTN+ mitophagy events were dual-labeled (**Figure S2B-C**). In contrast, when sequentially pulsed, 87.5% of OPTN+ mitophagy events were labeled exclusively by the first SNAP pulse ligand (**Figure S2D-F**). These results demonstrate that the SNAP pulse-chase assay can spectrally isolate mitochondrial fragments undergoing OPTN-dependent mitophagy.

To extend this approach to longer timescales, we added blocking and recovery steps to further separate the first and second SNAP ligand pulses temporally. After a 6-hour treatment and the first SNAP ligand pulse, we returned the neurons to control media. Neurons were then labeled with SNAP Block, a non-fluorescent ligand that saturates any remaining SNAP binding sites. The cultures recovered for 24 or 72 hours before labeling with the second, spectrally distinct SNAP ligand and LysoT (**Figure 4C**). We visualized spectrally distinct ‘Old’ and ‘Young’ mitochondrial populations within the dual-labeled network (**Figure 4D**). We identified OPTN+ ‘Old’ mitochondrial fragments and quantified the percentage that were within a lysosome (LysoT+; **Figure 4E**). While we observed no differences between control and AOF conditions, the percentage of OPTN+ ‘Old’ mitophagy events that colocalized with LysoT increased from 24 to 72 hours in AOF conditions (**Figure 4F**). Overall, these data demonstrate that OPTN-mediated mitophagy is a slow process occurring over days in primary hippocampal neurons and that, despite delayed autophagosome initiation, damaged mitochondria are ultimately degraded by lysosomes.

Given that autophagosome initiation and formation emerged as rate-limiting in OPTN-mediated neuronal mitophagy, we asked whether regulators of autophagy and lysosome function were altered. High levels of mitochondrial stress cause dose-dependent degradation of Myotubularin-related phosphatases (MTMR)2 and MTMR5, and Rubicon (Basak and Holzbaur, 2025).

MTMR2 and MTMR5 inhibit autophagosome biogenesis, while Rubicon blocks lysosome maturation and function (Basak and Holzbaur, 2025; Chua et al., 2022). We examined expression levels at 6, 24, and 72 hours and expected these negative regulators to decrease at 72 hours, consistent with the timing of mitochondrial degradation in our pulse-chase analysis.

However, at the later 72-hour time point, when we identified colocalization of OPTN+ mitochondria with LysoT, we still did not observe decreases in MTMR2, MTMR5, or Rubicon (**Figure S2G-J**). It is possible that we could not detect small changes in protein abundance using immunoblotting, or alternative constraints may limit OPTN-dependent mitophagy prior to 72 hours.

### Local ROS is elevated at OPTN+ mitophagy events

Since autophagosome initiation and engulfment are specifically impaired at OPTN+ mitophagy events, we hypothesized that local mitochondrial features, such as the bioenergetic state of damaged mitochondria, underlie this selective stalling. We examined local ATP and ROS levels at OPTN+ mitophagy events compared with the neuron or mitochondrial network, respectively. Mitochondria produce ATP through oxidative phosphorylation, and damaged mitochondria produce less ATP (Rangaraju et al., 2014). ATP is also important for autophagosome formation. For example, unc-51 like kinase 1 (ULK1), a key autophagy protein, uses energy to initiate autophagosome formation (Kim et al., 2011). We predicted that local ATP depletion at OPTN+ mitophagy events hinders efficient autophagosome initiation and engulfment. To test this, we used the ratiometric ATP sensor QUantitative Evaluator of cellular ENergy 37C (QUEEN-37C). The GFP chromophore in the ATP-bound state (400 nm) shifts to a higher excitation wavelength in the ATP-free state (494 nm) (Yaginuma and Okada, 2021). Intracellular ATP levels were visualized and quantified using 400/494 nm fluorescence intensity or the QUEEN Ratio. Neurons were transfected with Mito-SNAP, Halo-OPTN, and QUEEN-37C. Using potent ATP synthesis inhibitors, 2-deoxy-D-glucose (2-DG) and OA, we observed a significant reduction in the QUEEN ratio compared to controls (**Figure 5A-B**). We then compared neuronal ATP levels in the soma with those at OPTN+ mitophagy events after 6 hours of AOF treatment (**Figure 5C-E**). We found no differences in the QUEEN ratio between neuronal soma and OPTN+ mitophagy events.

**Figure 5:**
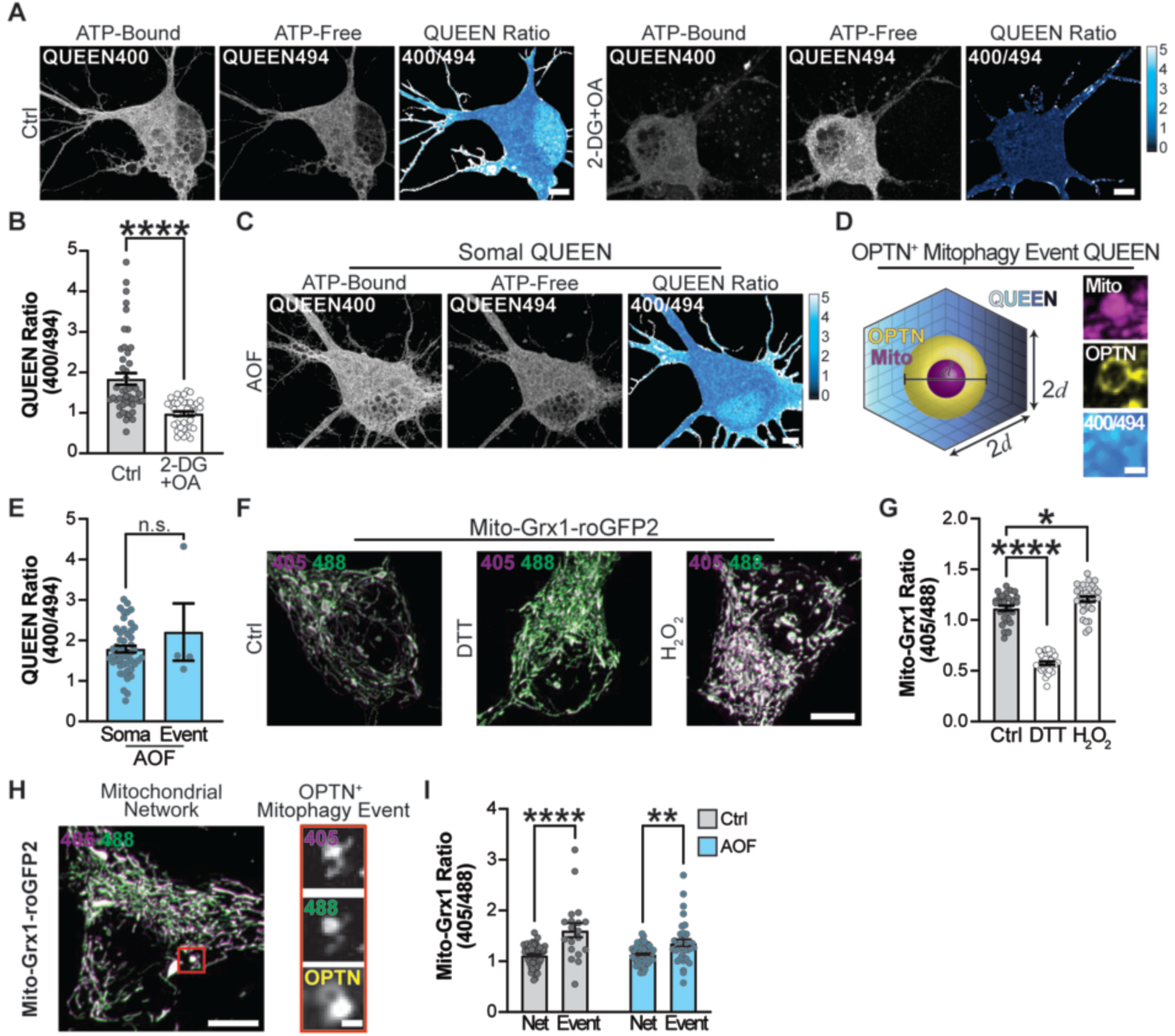
Mitochondrial ROS, but not ATP, is significantly higher at OPTN+ mitophagy events compared to the mitochondrial network. (**A**) Representative images of the QUEEN-37C in either control or 2-deoxyglucose + Oligomycin A (2-DG+OA). The QUEEN ratio is displayed in a LUT. Scale bar, 5 µm. (**B**) Quantification of the QUEEN ratio in the soma. Mean ± SEM; *n* = 39-46 neurons from 5 biological replicates; 7 DIV. ****, *p* < 0.0001 by unpaired Mann-Whitney test. (**C**) Representative images of the ATP sensor, QUEEN-37C, in the soma of primary hippocampal neurons. The QUEEN ratio is displayed in a LUT. Scale bar, 5 µm. (**D**) Schematic (left) and representative images (right) of the QUEEN ratio around an OPTN+ mitophagy event. The QUEEN ratio was determined within a 3D volume twice the diameter of the OPTN+ mitophagy event. Scale bar, 1 µm. (**E**) Quantification of the QUEEN ratio within the soma and at OPTN+ mitophagy events. Mean ± SEM; *n* = 4 events in 48 neurons from 5 biological replicates; 7 DIV. n.s., not significant by Welch’s t-test. (**F**) Representative images of Mito-Grx1-roGFP in either control, DTT, or H_2_O_2_ conditions. Scale bar, 5 µm. (**G**) Quantification of the Mito-Grx1 Ratio within the mitochondrial network. Mean ± SEM; *n* = 30-31 neurons from 3 biological replicates; 8-9 DIV. *, *p* < 0.05; ****, *p* < 0.0001 by One-Way ANOVA with Dunnett’s multiple comparisons test. (**H-I**) Representative images (H) and quantification (I) of Mito-Grx1-roGFP2 in the mitochondrial network and an OPTN+ mitophagy event. Scale bar: 5 µm (left); 1 µm (right). Mean ± SEM; *n* = 19-36 events from 6 plates per condition, which includes 56-58 neurons across 3 biological replicates; 8-9 DIV. **, *p* < 0.01; ****, *p* < 0.0001 by Two-Way ANOVA with Šídák’s multiple comparisons test. Net, network.

Next, we examined local ROS at OPTN+ mitochondrial fragments as a feature that could restrict autophagosome initiation and engulfment. While elevated ROS triggers OPTN-mediated mitophagy, it can also inhibit autophagy proteins and autophagosome formation (Zheng et al., 2020). To assess ROS, we used Mito-Grx1-roGFP2, which contains a mitochondrial-targeted glutaredoxin and a redox-sensitive roGFP2. This fluorophore excites at 488 nm under reducing conditions but shifts to 405 nm in oxidized environments. We therefore visualized and measured intracellular ROS as the 405/488 nm fluorescence intensity ratio (Mito-Grx1 Ratio). Neurons were transfected with Halo-OPTN and Mito-Grx1-roGFP2. We initially validated the reporter’s sensitivity using dithiothreitol (DTT) or hydrogen peroxide (H_2_O_2_), which reduce or oxidize the environment, respectively, and significantly shift the Mito-Grx1 Ratio (**Figure 5F-G**). The Mito-Grx1 Ratio was significantly higher at OPTN+ mitophagy events compared to the neuronal mitochondrial network in both control and AOF conditions (**Figure 5H-I**). Overall, these data show that elevated ROS, not ATP, distinguishes OPTN+ mitochondria and suggest that ROS may affect autophagosome initiation and engulfment at these sites.

### Scavenging mitochondrial ROS accelerates OPTN-mediated mitophagy progression in neurons

We next assessed whether selective ROS depletion could promote mitochondrial turnover by accelerating autophagosome initiation and engulfment. We scavenged ROS using N-acetylcysteine (NAC), an antioxidant and glutathione precursor, or EUK-134, a superoxide dismutase (SOD) mimetic that preferentially targets mitochondrial superoxide. We preferred this antioxidant treatment to overexpression because it allowed us to first induce mild ROS via AOF to initiate OPTN-mediated mitophagy, then immediately scavenge ROS to observe the downstream effects.

First, we induced mitophagy with AOF for 6 hours, then scavenged ROS for 24 hours with NAC or DMSO (**Figure 6A**). We measured mitochondrial ROS levels using MitoSOX, a dye that detects mitochondrial superoxide. We found a significant decrease in the normalized MitoSOX intensity in NAC-treated neurons compared to controls (**Figure 6B-C**), indicating lower ROS levels. We then used LysoT to visualize lysosomal degradation of OPTN+ mitophagy events. Neurons were transfected with Mito-DsRed2, Halo-OPTN, and labeled with LysoT. NAC treatment did not significantly alter the percentage of neurons undergoing OPTN-mediated mitophagy (**Figure 6D**) or somal mitochondrial network volume (**Figure S3A-B**) compared with controls. Interestingly, NAC-mediated ROS scavenging significantly increased LysoT+ OPTN-mediated mitophagy events (**Figure 6E-F**).

**Figure 6:**
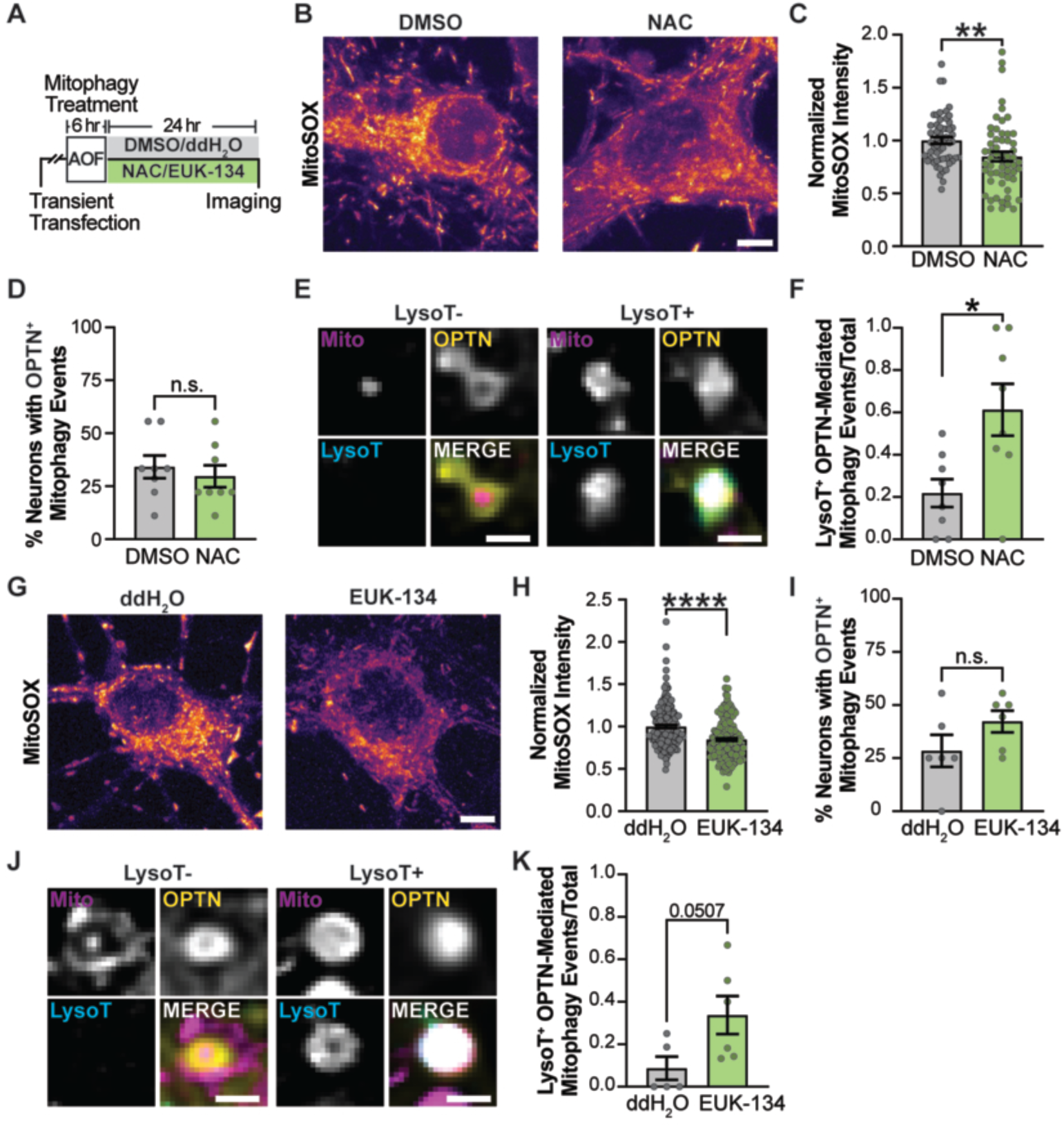
Scavenging intracellular or mitochondrial ROS accelerates OPTN-mediated mitophagy. (**A**) Schematic of the NAC or EUK-134 experimental timeline. Briefly, a 6-hour AOF treatment induces OPTN-mediated mitophagy, followed by 24 hours in DMSO, ddH_2_O, NAC, or EUK-134 to scavenge ROS. (**B-C**) Representative images (B) and quantification (C) of the MitoSOX fluorescence intensity in DMSO- and NAC-treated neurons. Mean ± SEM; *n* = 56-57 neurons from 3 biological replicates; 8 DIV. **, *p* < 0.01 by unpaired Mann-Whitney test. (**D**) Quantification of the percentage of DMSO- and NAC-treated neurons with OPTN+ mitophagy events. Mean ± SEM; *n* = 44-55 events from 8 plates per condition, which includes 70 neurons across 4 biological replicates; 8 DIV. n.s., not significant by unpaired t-test. (**E-F**) Representative image (E) and quantification (F) of the fraction of LysoT+ OPTN-mediated mitophagy events in DMSO- and NAC-treated neurons. Mean ± SEM; *n* = 44-55 events from 8 plates per condition, which includes 70 neurons across 4 biological replicates; 8 DIV. *, *p* < 0.05 by unpaired t-test. (**G-H**) Representative images (G) and quantification (H) of the MitoSOX fluorescence intensity in ddH_2_O and EUK-134-treated neurons. Mean ± SEM; *n* = 183-184 neurons from 4 biological replicates; 8 DIV. ****, *p* < 0.0001 by unpaired Mann-Whitney test. (**I**) Quantification of the percentage of ddH_2_O and EUK-134-treated neurons with OPTN+ mitophagy events. Mean ± SEM; *n* =47-64 events from 6 plates per condition, which includes 51-52 neurons across 3 biological replicates; 8 DIV. n.s., not significant by unpaired t-test. (**J-K**) Representative image (J) and quantification (K) of the fraction of LysoT+ OPTN-mediated mitophagy events in ddH_2_O and EUK-134-treated neurons. Mean ± SEM; *n* =47-64 events from 6 plates per condition, which includes 51-52 neurons across 3 biological replicates; 8 DIV. *p* = 0.0507 by unpaired t-test.

We repeated these experiments with EUK-134, a mitochondrial-targeted scavenger. EUK-134 efficiently scavenged mitochondrial ROS, significantly decreasing MitoSOX intensity compared with the control (**Figure 6G-H**). We again observed no differences in the percentage of neurons with OPTN+ mitophagy events (**Figure 6I**) or somal mitochondrial network volume (**Figure S3C-D)** in control and EUK-134-treated neurons. Similar to NAC, ROS scavenging with EUK-134 increased LysoT+ OPTN-mediated mitophagy events (**Figure 6J-K**). Together, these data indicate that mild oxidative stress promotes OPTN-mediated mitophagy, while scavenging elevated local ROS at damaged mitochondria is sufficient to overcome the rate-limiting step of autophagosome initiation and engulfment and promote progression through subsequent mitophagy steps within 24 hours.

### Partial nutrient deprivation induces autophagy and accelerates OPTN-dependent mitophagy

Given that autophagosome initiation and formation are rate-limiting, we next asked whether alterations to autophagy alone could affect the time course of OPTN-mediated mitophagy. We used neuronal starvation to stimulate autophagy by employing a partial nutrient-deprivation paradigm, where neurons were incubated with equal volumes of neuron culture media and Hanks’ Balanced Salt Solution (HBSS; (Wojnacki et al., 2020; Wojnacki et al., 2021)). We treated neurons with AOF for 6 hours to induce mitophagy, then incubated them in either control or HBSS for 24 hours (**Figure 7A**).

**Figure 7:**
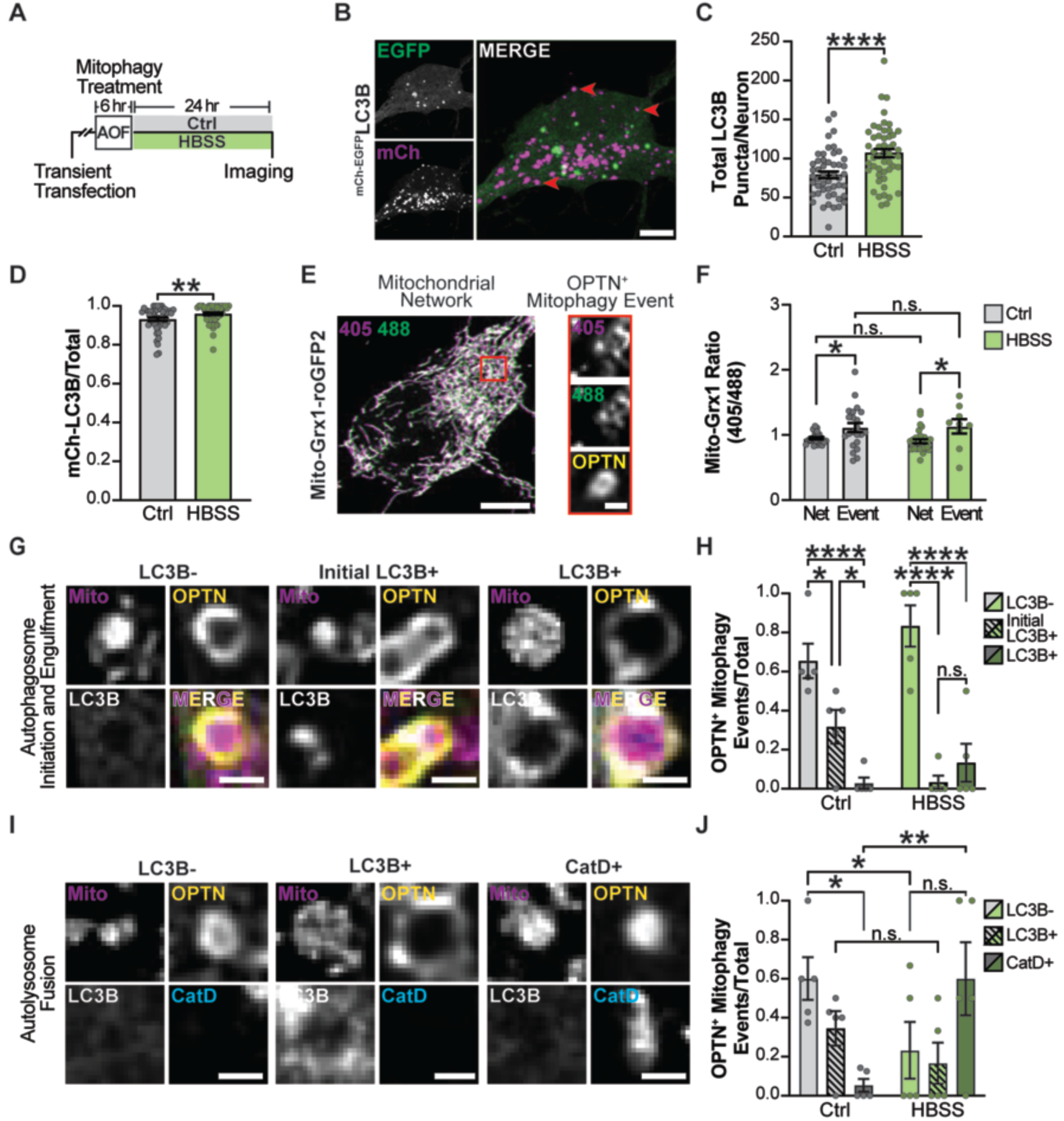
Neuronal starvation accelerates OPTN-mediated mitophagy. (**A**) Schematic illustrating the neuronal starvation experimental timeline. Briefly, a 6-hour AOF treatment induces OPTN-mediated mitophagy, followed by 24 hours in either control or HBSS to induce partial nutrient deprivation. (**B**) Representative image of autophagosomes in primary neurons labeled with mCherry-EGFP-LC3B. Red arrows highlight acidified autophagosomes. Scale bar, 5 µm. (**C**) Quantification of total LC3B puncta per neuron. Mean ± SEM; *n* = 49-50 neurons from 3 biological replicates; 9 DIV. ****, *p* < 0.0001 by unpaired t-test. (**D**) Quantification of the fraction of acidified autophagosomes (mCherry+ LC3B). Mean ± SEM; *n* = 49-50 neurons from 3 biological replicates; 9 DIV. **, *p* < 0.01 by unpaired Mann-Whitney test. (**E-F**) Representative images (E) and quantification (F) of Mito-Grx1-roGFP2 within the mitochondrial network and at OPTN+ mitophagy events. Scale bar, 5 µm (Left); 1 µm (Right). Mean ± SEM; *n* = 9-22 events from 3 plates per condition, which includes 24-28 neurons across 3 biological replicates; 9 DIV. n.s., not significant; *, *p* < 0.05; Two-Way ANOVA with Šídák’s multiple comparisons test. Net, network. (**G-H**) Representative images (G) and quantification (H) of OPTN+ mitophagy events that are LC3B-, Initial LC3B+, and LC3B+ following control or HBSS conditions. Scale bar, 1 µm. Mean ± SEM; *n* = 17-27 events from 5 plates per condition, which includes 43-45 neurons across 5 biological replicates; 9 DIV. n.s., not significant; *, *p* < 0.05; ****, *p* < 0.0001 by Two-Way ANOVA with Tukey’s multiple comparisons test. (**I-J**) Representative images (I) and quantification (J) of the fraction of OPTN+ mitophagy events that are LC3B-, LC3B+, or CatD+ following control or HBSS conditions. Scale bar, 1 µm. Mean ± SEM; *n* = 17-27 events from 5 plates per condition, which includes 43-45 neurons across 5 biological replicates; 9 DIV. n.s., not significant; *, *p* < 0.05; **, *p* < 0.01 by Two-Way ANOVA with Tukey’s multiple comparisons test.

To confirm that our partial nutrient-deprivation paradigm induced autophagy, we measured ribosomal protein S6 and LC3B levels in neuronal lysates. The negative autophagy regulator mechanistic target of rapamycin complex 1 (mTORC1) phosphorylates S6 kinase, which in turn phosphorylates the ribosomal protein S6. Therefore, phosphorylated S6 (pS6) serves as a readout of mTORC1 activity (Biever et al., 2015). Partial nutrient deprivation significantly reduced pS6 levels compared to control, while total S6 levels remained constant, and increased the LC3B-II: LC3B-I ratio (**Figure S4A-E**). We also visualized autophagosome abundance in neurons transfected with mCherry-EGFP-LC3B (**Figure 7B**). We observed a significant increase in both the total number of neuronal autophagosomes and the fraction of acidified (mCherry+) autophagosomes under HBSS conditions compared with the control (**Figure 7C-D**). These results indicate that mTORC1 signaling is inhibited, enhancing LC3B lipidation and autophagosome formation.

The response to nutrient deprivation appears context-dependent, as reports conflict on whether it consistently stimulates autophagosome formation in primary neuron cultures (Maday and Holzbaur, 2016; Young et al., 2009). So, we performed similar experiments using a previously reported 4-hour deprivation paradigm. Consistent with previous findings (Maday and Holzbaur, 2016), we observed significantly reduced pS6 levels but no change in the LC3B-II:LC3B-I ratio compared with the control (**Figure S4F-J**). Thus, the duration of nutrient deprivation appears to be an important determinant for the autophagic response.

Importantly, we confirmed that HBSS treatment did not affect the mitochondrial network or mitophagy but instead altered only autophagy. We treated neurons with control or AOF for 6 hours, then incubated them in control or HBSS for 24 hours (**Figure S5A**). We found no differences in mitochondrial volume, the percent of neurons with OPTN+ mitophagy events, or the number of events per neuron, regardless of the initial treatment (**Figure S5B-E**). Notably, we consistently observed reduced TMRE intensity and an increased Mito-Grx1 Ratio at OPTN+ mitophagy events compared to the mitochondrial network, indicating that HBSS preserves increased local ROS at these damaged mitochondria (**Figures 7E-F and S5F-I**). Together, these data validate our partial nutrient-deprivation paradigm as an effective method for broadly inducing neuronal autophagy without affecting OPTN-mediated mitophagy.

Next, we assessed whether nutrient deprivation-induced autophagy could increase autophagosome initiation and formation to accelerate OPTN-mediated neuronal mitophagy progression. Neurons were transfected with Mito-SNAP, Halo-OPTN, EGFP-LC3B, and CatD-RFP. We first visualized OPTN+ mitophagy events that were LC3B-, Initial LC3B+, and LC3B+ (**Figure 7G**). As we observed previously after 6 hours of treatment, most OPTN+ mitophagy events appeared to be in an LC3B-state when visualized with EGFP-LC3B in control and HBSS conditions (**Figure 7H**). Notably, HBSS had more LC3B-events than control. To determine whether this difference led to more OPTN+ mitophagy events progressing through the pathway to lysosomal fusion, we visualized OPTN+ mitophagy events and categorized them as outside a lysosome (CatD-) or within a lysosome (CatD+; **Figure 7I**). Consistent with the 24-hour SNAP pulse-chase analysis (**Figure 4F**), we observed that only a small fraction of OPTN+ mitochondria colocalized with CatD in the neurons incubated in control conditions. However, the fraction of CatD+ events increased significantly under HBSS conditions (**Figure 7J**), suggesting that OPTN+ damaged mitochondria are efficiently engulfed by autophagosomes and fuse with lysosomes for degradation. Overall, these findings indicate that increasing autophagic capacity via HBSS can overcome the ROS-associated delay and accelerate the progression of OPTN-mediated mitophagy in hippocampal neurons.

## DISCUSSION

Here, we combined multi-channel live-cell imaging with a physiologically relevant AOF paradigm to investigate the temporal dynamics of OPTN-dependent mitophagy, which is downstream of PINK1 and Parkin, in primary hippocampal neurons. By temporally resolving the individual steps of OPTN-mediated mitophagy, we identified autophagosome initiation and engulfment as rate-limiting, resulting in slow turnover that occurs over days. Inefficient autophagosome formation was observed at OPTN+ mitochondria, which displayed locally elevated ROS. Importantly, scavenging local mitochondrial ROS accelerated OPTN-facilitated mitophagy. Stimulating autophagy via neuronal starvation also accelerated OPTN-mediated mitophagy, indicating that pathway kinetics in neurons are constrained by autophagosome initiation and engulfment. Together, these findings provide mechanistic insights into slow pathway dynamics of OPTN-dependent mitophagy in primary hippocampal neurons (**Figure 8**).

**Figure 8:**
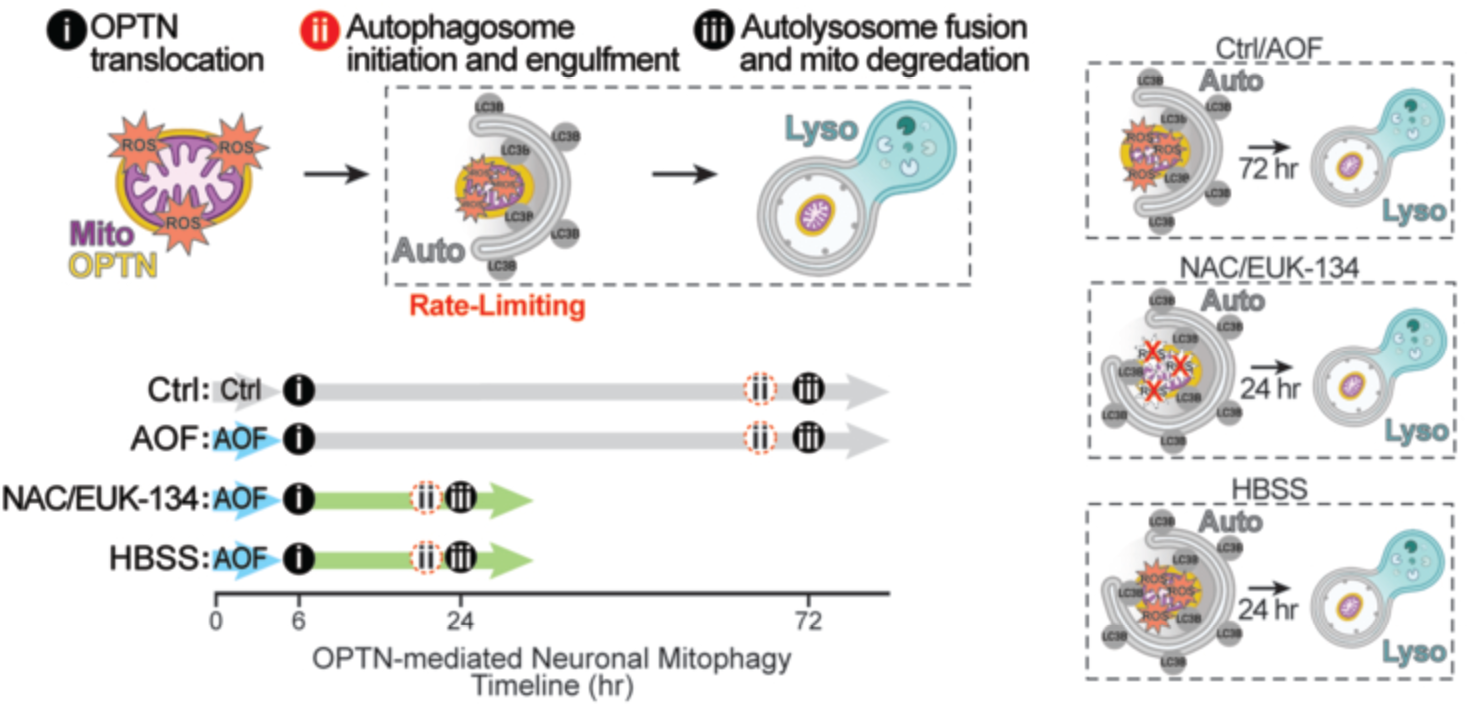
Summary schematic of the OPTN-mediated neuronal mitophagy timeline. **i**, OPTN translocates to damaged mitochondrial fragments. **ii**, An autophagosome initiates and engulfs an OPTN+ mitochondrion. **iii**, The formed autophagosome fuses with a lysosome, and the damaged mitochondria are degraded. Under control and AOF conditions, autophagosome initiation and engulfment are rate-limiting, delaying lysosomal fusion until 72 hours after the initial insult. In contrast, lysosomal fusion of damaged mitochondria rapidly occurs 24 hours after OPTN-mediated mitophagy induction due to ROS scavenging via NAC or EUK-134, or global autophagy induction via HBSS. The exact kinetics of autophagosome formation remain unresolved (ii, dashed, bottom timeline), but appear to occur prior to rapid lysosome fusion.

We identified autophagosome initiation and engulfment as rate-limiting in OPTN-mediated neuronal mitophagy (**Figure 2**). This conclusion differs from our previous report, where we observed efficient autophagosome fusion and reported lysosomal acidification as the limiting factor (Evans and Holzbaur, 2020a; Evans and Holzbaur, 2020b). We attribute this discrepancy to differences in the OPTN events quantified. Here, we broadened our inclusion criteria to capture both OPTN+ rings and puncta, whereas we previously used only OPTN+ rings. Interestingly, this change significantly increased the number of OPTN+ mitophagy events that were LC3B- and had not yet undergone autophagosome engulfment. These differences now provide insights into OPTN organization during mitophagy. We propose that OPTN may initially translocate to damaged mitochondria as puncta and then reorganize into ring-like structures as the pathway progresses to support LC3B recruitment. Consistent with this model, LC3B was previously visualized first as puncta that redistributed into rings that colocalized with Parkin. Over time, both Parkin and LC3B transitioned to puncta that overlapped with damaged mitochondria (Li et al., 2021). Alternatively, OPTN puncta and rings may represent separate populations, and LC3B may be preferentially recruited to or retained at OPTN+ rings. Recent work shows that OPTN forms dynamic, sheet-like phase-separated condensates and that OPTN fluidity alters autophagic membrane recruitment (Yang et al., 2024).

Our data place the temporal dynamics of OPTN-dependent neuronal mitophagy on the order of days (**Figure 4**). Furthermore, AOF increased the frequency of OPTN+ mitophagy events but did not alter kinetics, suggesting the time course is intrinsic to neurons. However, currently there is no consensus on neuronal mitophagy kinetics, and the mechanism underlying these discrepancies remains unclear. Differences likely reflect the type or severity of mitochondrial stress, the neuronal compartment examined, or variability across neuronal populations. Our findings are consistent with prior work, where sustained ROS burden ranging from mild oxidative stress paradigms (Evans and Holzbaur, 2020a; Li et al., 2021) to more global mitochondrial depolarization (Cai et al., 2012; Lin et al., 2017) may stall or delay the time course of mitophagy in the soma. Thus, while neurons can recognize, remodel, and sequester damaged mitochondria, downstream mechanisms to fully complete mitophagy may be delayed in stress-dependent contexts. On the other hand, another study reported rapid mitophagy kinetics following an acute high dose of AA (Ashrafi et al., 2014). However, this study focused exclusively on axonal mitochondria and captured events immediately after insult. Our data support a model where OPTN-facilitated mitophagy is intrinsically slow in the soma of hippocampal neurons, and these kinetics are unaltered under prolonged, mild physiological stress.

We found that neurons treated with AOF had higher intracellular ROS compared to control (**Figure 1B-C**). However, local mitochondrial ROS levels at OPTN+ mitophagy events were elevated relative to those in the neuronal mitochondrial network in both control and AOF conditions. We found that scavenging mitochondrial ROS had no effect on OPTN-mediated mitophagy initiation, but accelerated mitophagy progression through lysosomal fusion (**Figure 6**). This supports a model that pathway kinetics are determined by the local redox environment at damaged mitochondria rather than by global cellular ROS levels. Mechanistically, local oxidative stress could alter the activity of autophagy-related proteins that drive autophagosome formation (Zheng et al., 2020). Alternatively, local ROS could modify OPTN fluidity to constrain interactions with the autophagic machinery. Future research will clarify the link between increased ROS and impaired autophagosome formation. Furthermore, scavenging ROS could serve a therapeutic function by mitigating their deleterious effects. Indeed, ROS scavenging with mitochondrially-targeted MitoTEMPO rescued mitochondrial defects in amyloid beta-treated hippocampal neurons (Hu and Li, 2016). These findings argue that mitochondrial and neuronal health are highly dependent on redox states.

We found that partial nutrient deprivation via HBSS induced autophagy, which was sufficient to increase the progression of OPTN-mediated mitophagy in neurons (**Figure 7**). Interestingly, we still observed elevated ROS and decreased TMRE fluorescence intensity at OPTN+ mitophagy events, but no change in either the percentage of neurons with OPTN+ mitophagy events or the number of events per neuron (**Figure S5**). Thus, increasing autophagosome formation is sufficient to bypass the inhibitory effects of local ROS at delayed OPTN+ mitophagy events and promote mitophagy pathway progression. ROS is thought to contribute to autophagy activation, which helps mitigate ROS levels by engulfing aged organelles, such as damaged mitochondria (Li et al., 2015). In HeLa cells, starvation increased superoxide levels to initiate autophagosome formation (Chen et al., 2009). However, in our system, we did not observe differences in the Mito-Grx1 Ratio between control and HBSS conditions (**Figure 7E-F**). Future work is needed to determine how local ROS at OPTN+ mitophagy events directly interacts with autophagosome formation within our system.

Though seemingly counterintuitive, these intrinsically slow kinetics may also serve an adaptive role. Given the high dependence of neuronal function on mitochondria, preserving dysfunctional mitochondria may be beneficial. Damaged mitochondrial components could be rapidly catabolized into reusable nutrients, such as amino acids, fatty acids, and sugars, to drive neuronal responses to stress (Kaur and Debnath, 2015). In addition, mitophagy may function as a secondary mitochondrial quality control mechanism preceded by a faster repair pathway. Mitochondrial-derived vesicles (MDVs) can repair damaged mitochondria by budding from them and removing damaged proteins and lipids, thereby preserving the parent organelle’s health (McLelland et al., 2014). For example, MDVs positive for syntaphilin (SNPH), a mitochondrial brake, form to remove SNPH from mitochondria and favor the transport of stressed mitochondria out of axons (Lin et al., 2017). Only after irreversible mitochondrial damage may neurons upregulate OPTN-dependent mitophagy to degrade dysfunctional mitochondria.

Mitochondrial function is a key determinant of neuronal health, and deficits in neuronal mitochondrial quality control are common across neurodegenerative diseases. Our results imply that intrinsically slow OPTN-dependent mitophagy kinetics cause damaged mitochondria to persist and accumulate, potentially increasing neuronal vulnerability to age- and disease-associated factors, such as decreased autophagosome formation, elevated oxidative damage, and altered lysosomal acidification (Lee et al., 2011; Lie and Nixon, 2019; Liguori et al., 2018; Stavoe et al., 2019). Consistent with this idea, the accumulation of damaged mitochondria in human hippocampi is linked to Alzheimer’s disease pathology (Fang et al., 2019). Mice lacking functional Parkin exhibit fragmented mitochondria, neuronal loss, and severe motor impairments, which are common in PD (Noda et al., 2020). Additionally, OPTN KO mice accumulate dysfunctional mitochondria in motor neurons, a feature seen in ALS (McCall et al., 2020). As these mitochondrial stress features appear to accumulate with age and disease, antioxidants like SOD remain relatively unchanged in brain tissue (Sohal et al., 1994). Together, these findings suggest that neuronal redox states are an important determinant for the temporal dynamics of OPTN-mediated mitophagy and that prolonged delays in mitochondrial turnover could contribute to the selective vulnerability of neurons in neurodegenerative disease.

## DATA AVAILABILITY

Data available upon request.

## ACKNOWLEDGEMENTS

We thank colleagues who generously provided reagents and members of the Evans Lab for their thoughtful feedback and comments on this manuscript. This work was supported by the Chan Zuckerberg Initiative BioHub (2022-253617), NIH T32 Training Grant (5T32GM142605-03 and 2T32GM142605-06), the Howard Hughes Medical Institute (HHMI) Hanna Gray Fellowship, McKnight Endowment for Neuroscience, and Duke Whitehead Scholars. Figures 1A, 2A, and 8 were created using BioRender (https://BioRender.com).

## AUTHOR CONTRIBUTIONS

Conceptualization, J.R.G. and C.S.E.; Methodology, all authors; Formal Analysis, all authors; Investigation, all authors; Writing – Original Draft, J.R.G. and C.S.E.; Writing – Review & Editing, all authors; Supervision, C.S.E.; Funding Acquisition, J.R.G. (training grant), L.D.P. (training grant), and C.S.E. (all other funding sources).

## MATERIALS AND METHODS

### Constructs and siRNA

Cathepsin D-RFP, Halo-OPTN, Mito-SNAP, Mito-TagBFP, Mito-EGFP-mCherry, and Untagged Parkin were kindly provided by Erika Holzbaur (University of Pennsylvania, Philadelphia). mCherry-EGFP-LC3B was provided by Terje Johansen (University of Tromso, Norway). EGFP-LC3B was provided by Tamotsu Yoshimori (Osaka University, Japan). Mito-DsRed2 was provided by Thomas Schwarz (Harvard Medical School, Boston). Mito-Grx1-roGFP2 (subcloned from pLPCX: mito-Grx1-roGFP2) was provided by Minna Roh-Johnson (University of Utah, Salt Lake City). pN1-QUE37C (QUEEN-37C) was a gift from Yasushi Okada (Addgene plasmid # 129318). ON-TARGET*Plus* Rat OPTN (246294) siRNA *SMARTpoo*l (L-097177-02-0005) was purchased from Horizon Discovery; target sequences include GCUAAAUAAUCAAGCUAUG, GGAGGAGAAAUCCAGGUUA, UGAAAGAACUCCUGGUCGA, and GAACAGCGCUGGACCCACA. All constructs were verified using whole plasmid sequencing (Plasmidsaurus).

### Reagents

Reagents used for experiments include, Antimycin A (Sigma-Aldrich, A8674-50MG or Cayman Chemicals, 34799), Oligomycin A (Sigma-Aldrich, 75351-5MG), B-27 Supplement minus antioxidants (ThermoFisher, 10889038), Bafilomycin A1 (Cell Signaling Technology, 54645S), CellROX Deep Red Reagent (Invitrogen, C10422), Cycloheximide (Sigma-Aldrich, 01810-1G), 2-Deoxy-D-Glucose (Sigma-Aldrich D6134-250MG), Dithiothreitol (Fisher BioReagents, BP172-5), anhydrous DMSO (Invitrogen, D12345), 1 M HEPES (Gibco, 15630-080), 10X HBSS (Gibco, 14185-052), cOmplete Mini EDTA-Free Protease Inhibitor Cocktail (Sigma-Aldrich, 11836170001), Halt Protease and Phosphatase Inhibitor Cocktail (100X) (ThermoFisher, 78440), Hydrogen peroxide solution (Sigma-Aldrich, H1009-5ML), LysoTracker Green DND-26 (ThermoFisher, L7526), MitoSOX Red (Invitrogen, M36008), N-acetylcysteine (Selleck Chemicals, S1623), EUK-134 (Sigma-Aldrich, SML0743), TMRE (tetramethylrhodamine ethyl ester, Ethyl Ester, Perchlorate; Life Technologies, T669), and Mycoplasma PCR Detection Kit (Applied Biological Materials, G238). SNAP and Halo constructs were labeled with the following ligands: SNAP-Cell 430 (New England Biolabs, S9109S), SNAP-Cell Block (New England Biolabs, S9106S), SNAP-JF549cp (Janelia Research Campus), HaloTag JF646 (Janelia Research Campus), and HaloTag JFX646 (Janelia Research Campus). We thank the Lavis lab and Open Chemistry team (Janelia) for the gift of JF and JFX dyes. For a list of key reagents, see the **Key Resources Table**.

### Primary hippocampal culture

Embryonic day 18 (E18) timed-pregnant Sprague Dawley rats were obtained from Charles River Laboratories. Hippocampi were isolated from all pups and dissociated to generate a cell suspension. Isolated hippocampal neurons were plated on 35 mm glass-bottom dishes (MatTek or CellVis) at 125,000 neurons for imaging analyses, 6-well plates at 125,000-250,000 neurons for immunoblotting analyses, or 96-well plates at 20,000 neurons for plate reader analyses. Imaging dishes and wells were precoated with 0.5 mg/mL poly-L-lysine (Sigma-Aldrich) for 18-24 hours prior to plating. Neurons were initially plated in MEM supplemented with 1 mM sodium pyruvate, 10% horse serum, and 33 mM D-glucose. After 3-5 hours, the media was replaced with Neurobasal (Gibco) supplemented with 100 units/ml penicillin, 100 mg/ml streptomycin, 2 mM GlutaMAX (Invitrogen), 33 mM D-glucose, and 2% B-27 (ThermoFisher), referred to as maintenance media, which was used to maintain the neurons for feeding and transfections. Neurons were maintained at 37 C in 5% CO_2_. AraC (1 µM) was added on DIV 1 to prevent the proliferation of non-neuronal cells. Neurons (6-8 DIV) were transfected (0.35-1.4 µg of total plasmid DNA) with Lipofectamine 2000 Transfection Reagent (ThermoFisher), and incubated for 18-24 hours. Plasmid DNA and siRNA (25 nM) mixtures were incubated for 36-48 hours.

### HeLa culture

Cells were maintained in Dulbecco’s modified Eagle medium (DMEM; Gibco) supplemented with 10% FBS (Cytiva Hyclone) and 1% GlutaMAX (Gibco) at 37 °C in 5% CO_2_. For transient transfection, cells were seeded on uncoated 35 mm glass-bottom dishes (MatTek) and transfected with a total of 2 µg of DNA using FuGene 6 (Promega) according to manufacturer’s instructions. Cells were incubated for 18 hr after transfection before subsequent treatment and imaging. We tested HeLa cells and confirmed they were free of mycoplasma contamination using the Mycoplasma PCR Detection Kit (ABM).

### Mitochondrial damage and treatments

Oxidative stress was induced with antioxidant-free (AOF) conditions or Antimycin A (AA; Sigma-Aldrich). For each 6-hour treatment, media was completely removed and replaced with either 1) AOF maintenance media, where B-27 Supplement minus antioxidants (ThermoFisher) replaced the standard B-27 supplement in the media, or 2) maintenance media containing 5 or 100 nM AA. For bafilomycin experiments, neurons were treated with 100 nM BafA1 (Cell Signaling Technology) throughout the 6-hour AOF treatment. For QUEEN-37C control experiments, neurons were treated with either control or 2-deoxy-glucose (2-DG; 30 mM) plus oligomycin A (OA; 1 µM) for 5 minutes, then imaged. For Mito-Grx1-roGFP2 control experiments, neurons were treated with either hydrogen peroxide (H_2_O_2_; 1 mM) or dithiothreitol (DTT; 1 mM) and then immediately imaged. For western blots, cycloheximide (10 µg/ml) was added for the duration of treatment. For antioxidant treatments, N-acetylcysteine (NAC; 500 µM) or EUK-134 (1 µM) were added after a 6-hour AOF treatment and neurons were incubated for 24 hours. For the neuronal starvation experiments, neurons were either maintained in control maintenance media or HBSS media (equivalent volumes of maintenance media and 1X HBSS [Gibco], supplemented with 10 mM HEPES [Gibco]) for 24 hours following the 6-hour control or AOF treatment.

Neurons were imaged in neuronal imaging media (Neurobasal medium, minus phenol red [Gibco] supplemented with 33 mM D-glucose, 2 mM GlutaMAX, 100 units/ml penicillin, 100 mg/ml streptomycin, and 2% B-27). To correspond with the experimental approach, the following treatments were added to the imaging media: AOF, the standard 2% B27 was replaced by 2% B-27 Supplement minus antioxidants; AA, imaging media contained 5 nM AA; BafA1, imaging media contained 100 nM BafA1 or equivalent volume of DMSO; HBSS, equivalent volumes of imaging media and 1X HBSS plus HEPES; QUEEN-37C controls, the imaging medium without glucose contained 30 mM 2-DG and 1 µM OA or equivalent volume of DMSO; Mito-Grx1-roGFP2 controls, 1 mM H_2_O_2_, 1 mM DTT, or equivalent volume of DMSO was added to the imaging medium at the time of imaging; NAC, imaging media contained 500 µM NAC; EUK-134, imaging media contained 1 µM EUK-134.

HeLa cells were treated with a combination of Antimycin A (0.5 or 4 µM) and Oligomycin A (0.5 or 4 µM) or an equivalent volume of ethanol for 2 hours. HeLa cells were imaged in phenol red-free DMEM supplemented with 10% FBS and 1% GlutaMAX (HeLa imaging media). To correspond with the experiment, the HeLa imaging media also contained 0.5 or 4 µM of AA/OA or vehicle control.

### SNAP and Halo labeling

One hour prior to live-cell imaging, SNAP (Cell 430, 2 µM; JF549cp, 50-100 nM) and Halo (JFX646 or JF646, 50-100 nM for neurons; JF646, 2.5 µM for HeLa cells) ligands were applied for 30 minutes, quickly washed twice, followed by a 30-minute washout, for a total of 1 hour. The neurons were then imaged for the following hour. For SNAP pulse-chase experiments, SNAP JF549cp (50 nM) and Halo JFX646 (50 nM) were added to AOF medium for 5 hours and 30 minutes, then washed twice quickly, followed by a 30-minute washout, for a total of 6 hours. Next, SNAP Block (0.5 µM) was added for 2 hours, followed by two quick washes, a 30-minute washout, and returned to control maintenance media for the recovery period. Finally, after 24 or 72 hours of recovery, SNAP Cell 430 (2 µM) was added for 30 minutes, followed by two quick washes, a 30-minute washout, and immediately imaged. The same procedure was used for the sequential SNAP pulse-chase, except that SNAP JF549 was added for a total of 6 hours without blocking or recovery steps, followed by SNAP Cell 430 for 1 hour. For the simultaneous SNAP pulse-chase experiment, SNAP JF549 and Cell 430 were co-incubated for a total of 6 hours without blocking and recovery steps. The same wash steps were performed after the addition of each ligand for sequential and simultaneous labeling.

### Dye labeling

To measure mitochondrial membrane potential, TMRE (2.5 nM) was loaded for 30 minutes, washed twice, and either immediately imaged with a confocal microscope or analyzed using a CLARIOstar^Plus^ plate reader (BMG LABTECH). The same TMRE concentration was also added to the imaging media for the plate reader analysis. To determine intracellular ROS, neurons were labeled with CellROX (5 µM) for 30 minutes, washed twice, and then immediately imaged. The same procedure was used for HeLa cells, except they were washed three times before immediate imaging. To assess lysosomal acidification, LysoTracker Green (25-50 nM for neurons; 50 nM for HeLa cells) was incubated for 30 minutes, then washed once before imaging. When applicable, TMRE, CellROX, and LysoTracker incubations were performed during the final 30 minutes of the 6-hour treatment (neurons), 2-hour treatment (HeLa cells), or the washout period of SNAP and Halo labeling. To measure mitochondrially produced ROS, MitoSOX Red (250 nM) was applied for 30 minutes, washed twice, and immediately imaged with a confocal microscope.

### Live-cell imaging

Images were acquired on a STELLARIS 8 Laser Scanning Confocal microscope (Leica) equipped with a 63x/1.4 NA oil-immersion objective using “Sequential Scanning” mode, 1024x1024 format (0.103 um/pixel), 300 nm step size (averaging 35-50 steps per image for neurons or 18-25 for HeLa cells), bidirectional scanning, 600 Hz scan speed, zoom factor of 1 or 1.75, line average of 2, and a pinhole size was 1 AU. For HeLa CellROX experiments, all parameters were constant; for neuron CellROX imaging experiments, we used “Stacked Sequential” mode with no line averaging. The microscope is housed in an environmental chamber maintained at 37 C and 5% CO_2,_ enabling live-cell imaging.

### Immunoblotting

Neurons were washed twice in PBS and lysed in RIPA buffer (50 mM Tris-HCl pH 7.4, 150 mM NaCl, 0.1% SDS, 0.5% deoxycholate, 0.1% Triton X-100, 1 mM PMSF, 1 mM DTT, and 1X complete protease inhibitor mixture) at 4 C for 30 minutes. For the 24 hour HBSS western blot samples, RIPA buffer was instead combined with Halt Protease and Phosphatase Inhibitor (ThermoFisher). All samples were centrifuged at 4 C for 10 minutes at 15,800 x g, and the supernatant was collected before determining the total protein concentration using a BCA assay. 15-20 µg of total protein were analyzed by SDS-PAGE and transferred to a PVDF Immobilon-FL (Millipore) membrane. Membranes were dried for 1 hour, rehydrated via methanol, and stained for total protein with REVERT Total Protein Stain (LI-COR). Once imaged, membranes were destained, blocked for 1 hour in Intercept Blocking Buffer TBS (Tris Buffered Saline; LI-COR), and incubated overnight with primary antibodies in Intercept Antibody Diluent (LI-COR) at 4 C. The membranes were then washed three times for 5 minutes in 1X TBS Washing Buffer (50 mM Tris-HCl pH 7.4, 274 mM NaCl, 5 mM KCl, 0.1% Tween-20) and incubated for 1 hour with secondary antibodies diluted in Intercept Antibody Diluent at room temperature. The membranes were washed three times for 5 minutes in 1X TBS Washing Buffer, followed by a final wash in 1X TBS, then imaged on the Odyssey XF (LI-COR) or ChemiDoc MP (Bio-Rad) imaging systems.

### Analysis

#### Image Processing

All images acquired through live confocal microscopy were deconvolved using the “Lightning Process” feature in the Leica Application Suite X (LAS X) Software. For CellROX, TMRE, Mito-Grx1-roGFP2, and MitoSOX Red experiments, fluorescence intensities were measured from unprocessed images. Images exhibiting frame shifts were registry corrected using an alignment macro in FIJI. Briefly, two slices were removed from the bottom of the misaligned z-stack and from the beginning of the aligned reference stacks. The misaligned channel was then translated +1 in both the *x* and *y* directions. The realigned z-stacks were then merged and used to generate a new, registry-corrected image file.

#### Image analysis

In neurons, OPTN+ mitophagy events were manually identified across single slices throughout the z-stack in FIJI, where they must traverse multiple z-planes to be considered. For HeLa cells, OPTN+ mitophagy events were manually identified from a single max-intensity projection generated from the corresponding z-planes in FIJI. Only mitophagy events that were clearly disconnected from the network and within an OPTN+ structure were analyzed. OPTN structures with a clear, hollow ring around the mitochondria were classified as “rings.” OPTN “puncta” were defined as solid, bright OPTN structures that either completely overlapped with the damaged mitochondrial fragment or extended beyond its boundary. OPTN+ mitophagy events were first identified based on our criteria without consideration of fluorescence signal from autophagosomes or lysosomes. We then returned to each event and manually identified the colocalization of associated markers with the OPTN+ mitophagy event. The fraction of OPTN-mediated mitophagy events positive and negative for structures of interest was calculated per imaging dish. Somal mitochondrial volumes were calculated using the intensity-based surface rendering feature in Imaris (Oxford Instruments) applied to the soma.

Surface detail was set to 0.001 µm, while surface threshold values were determined manually for each neuron. Surfaces smaller than the surface detail threshold were removed from the final volume calculation. Surfaces covering the background or mitochondria outside the soma were manually removed. The number of LC3B+ or CatD+ puncta was obtained using max projection of images in FIJI. To quantify acidified autophagosomes, the number of unacidified autophagosomes (mCherry-EGFP-LC3B puncta) was subtracted from the total number of autophagosomes (mCherry-LC3B puncta), then divided by the total number. To quantify general mitophagy using the dual-labeled Mito-EGFP-mCherry, the number of EGFP-/mCherry+ puncta was quantified using the Cell Counter feature in FIJI. For each neuron, the number of puncta was normalized to mitochondrial volume (calculated in Imaris).

#### CellROX, TMRE, Mito-Grx1-roGFP2, and MitoSOX Red quantification

For mitochondrial network quantification of CellROX, TMRE, Mito-Grx1-roGFP2, and MitoSOX Red, mean gray values were measured from max projections of unprocessed confocal images. In FIJI, the average of five areas across the soma was measured using a 1.34 x 1.34 or 2.3 x 2.3 µm square, or 2.2 x 2.2 µm for HeLa cells within the perinuclear area. We applied the same region size and analysis procedure to all experimental conditions and biological replicates. MitoSOX Red values were normalized to the average control fluorescence intensity within each biological replicate. For OPTN+ mitophagy event quantifications, TMRE and Mito-Grx1-roGFP2 fluorescence intensities were measured from single z-slices, where the event appeared most prominently. In FIJI, the mean gray values of five areas within each OPTN structure were measured using a 1 x 1 pixel square and then averaged. Mito-Grx1 Ratios were calculated by first averaging all five areas, then dividing the average fluorescence intensity values at 405 by 488. TMRE intensities (Figure 1F) were quantified using a CLARIOstar Plus plate reader (BMG LABTECH), averaging six technical replicates per biological replicate from three independent experiments.

#### QUEEN-37C quantification

Quantifications of the QUEEN ratio (400/494 nm) were obtained from deconvolved images. In FIJI, the average mean gray values of five areas within the soma were measured using a 1.2 x 1.2 µm box for the 400- and 494-channel images, then divided to generate a QUEEN ratio for each neuron. To generate QUEEN ratiometric images, max projections were produced for each channel, with background subtraction performed using the “Subtract” feature in FIJI. Representative ratiometric images were created using the “Image Calculator” with the operation (division) and 32-bit (float) selected to allow non-integer ratios in the final image. The LUT display was set to Cyan Hot, with the “Brightness & Contrast” scale set from 0 to 5. Each neuron was then traced using the polygon feature to display only the selected region.

#### Fluorescent immunoblotting quantification

Western blot bands were analyzed using Image Studio Software (LI-COR) or Image Lab (Bio-Rad). For each blot, total protein normalization factors were calculated by dividing each total protein lane by the lane with the highest signal. This approach was used instead of a housekeeping loading control and ensures consistent protein loading and transfer across the entire blot. The calculated total-protein normalization factors were used to normalize the bands of interest in each lane. To determine fold change, treatment band intensities were divided by control band intensities.

#### Line scan intensity profiles

The intensity profiles along the line of interest for an OPTN+ mitophagy events were generated using a line width of 1 in FIJI. The resulting values were normalized by dividing each by the maximum measured gray value for that channel.

Prism was used to calculate statistics and represent data graphically. Adobe Illustrator was used to assemble all figures, including images and graphs. Portions of schematics were generated using BioRender.

#### Declaration of generative AI

During the preparation of this work, the authors used ChatGPT (OpenAI) and Grammarly (Superhuman Platform Inc.) to improve readability and language; the authors reviewed and edited the content as needed after use. The authors take full responsibility for the content in this publication.

---

**Table 1:**
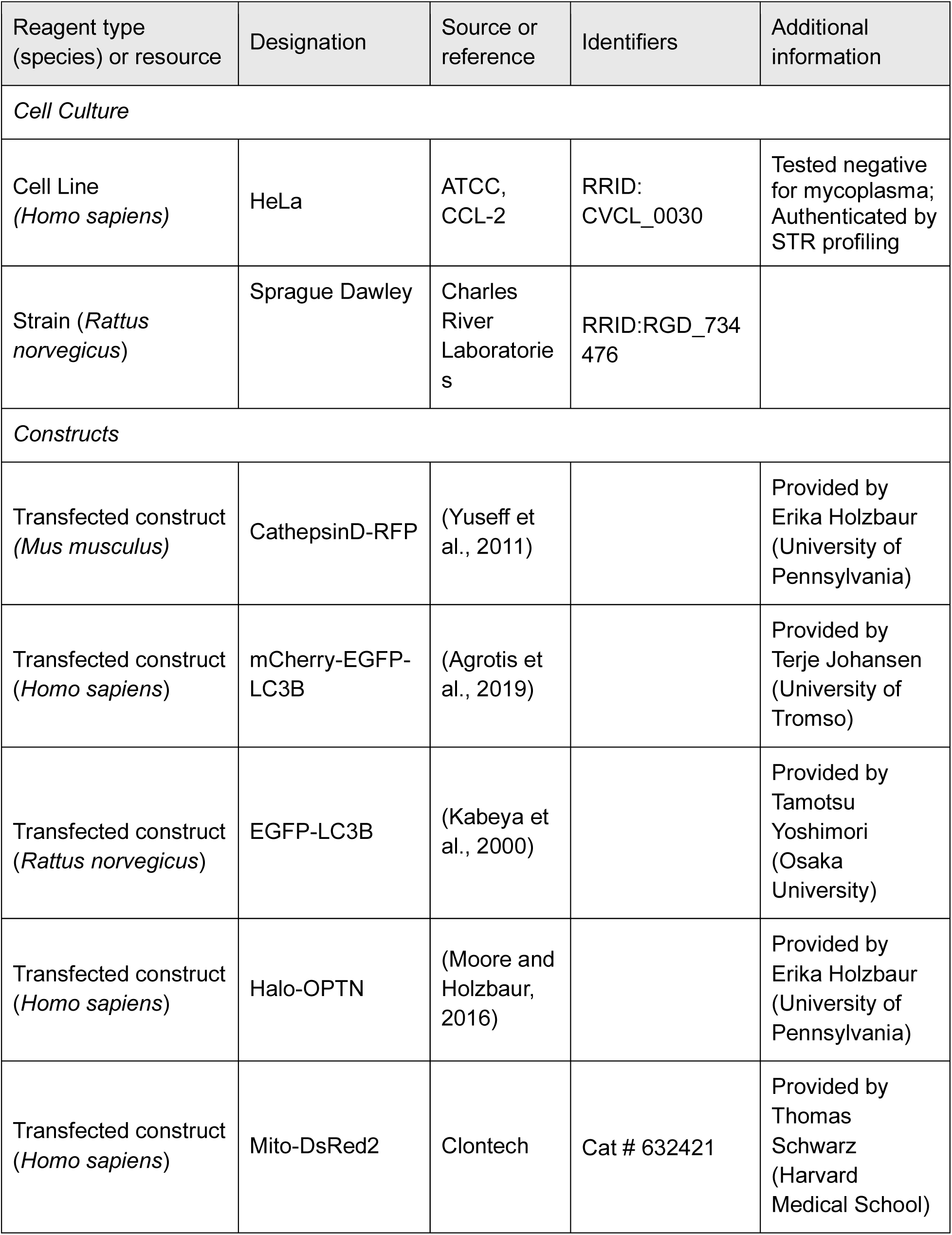
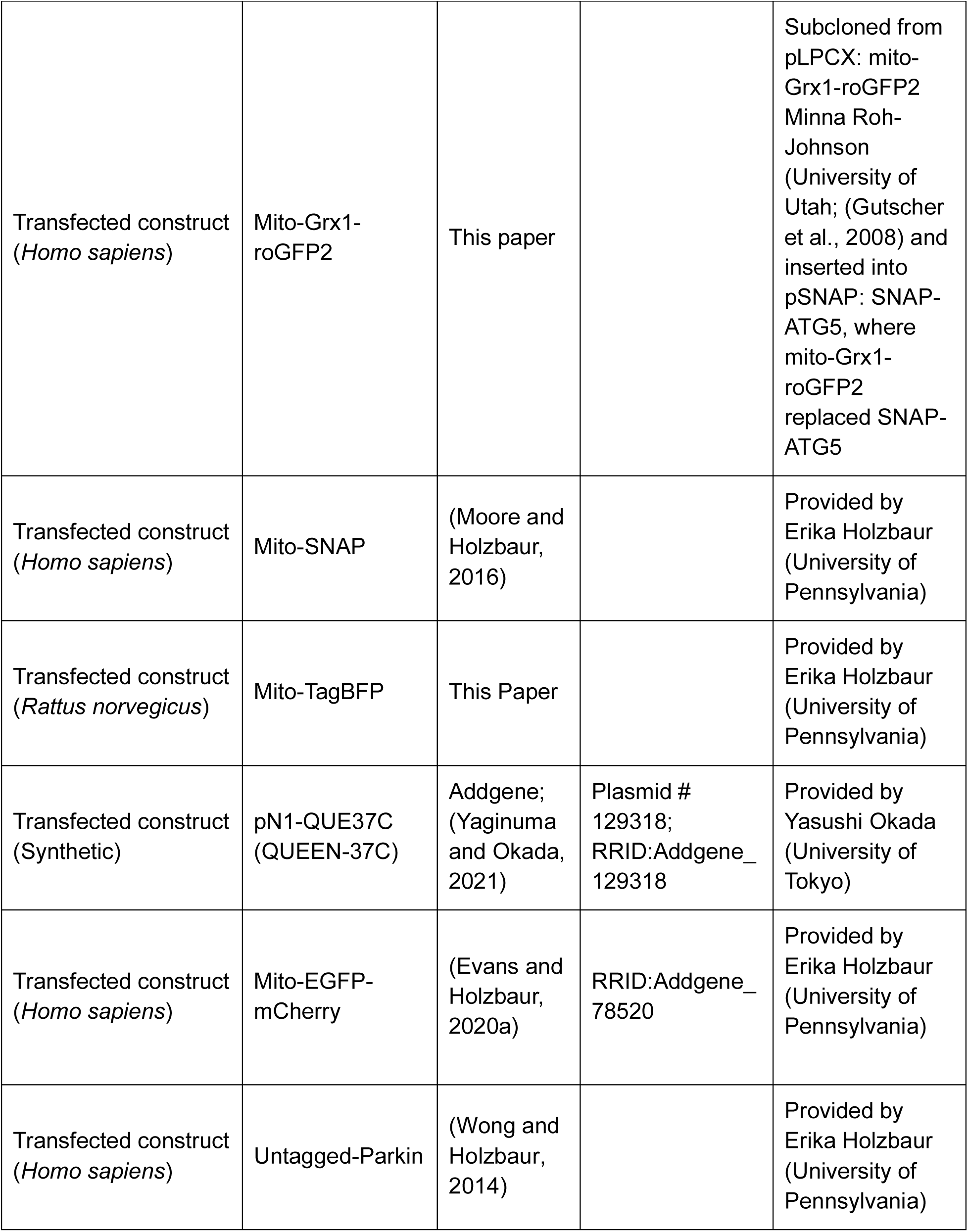
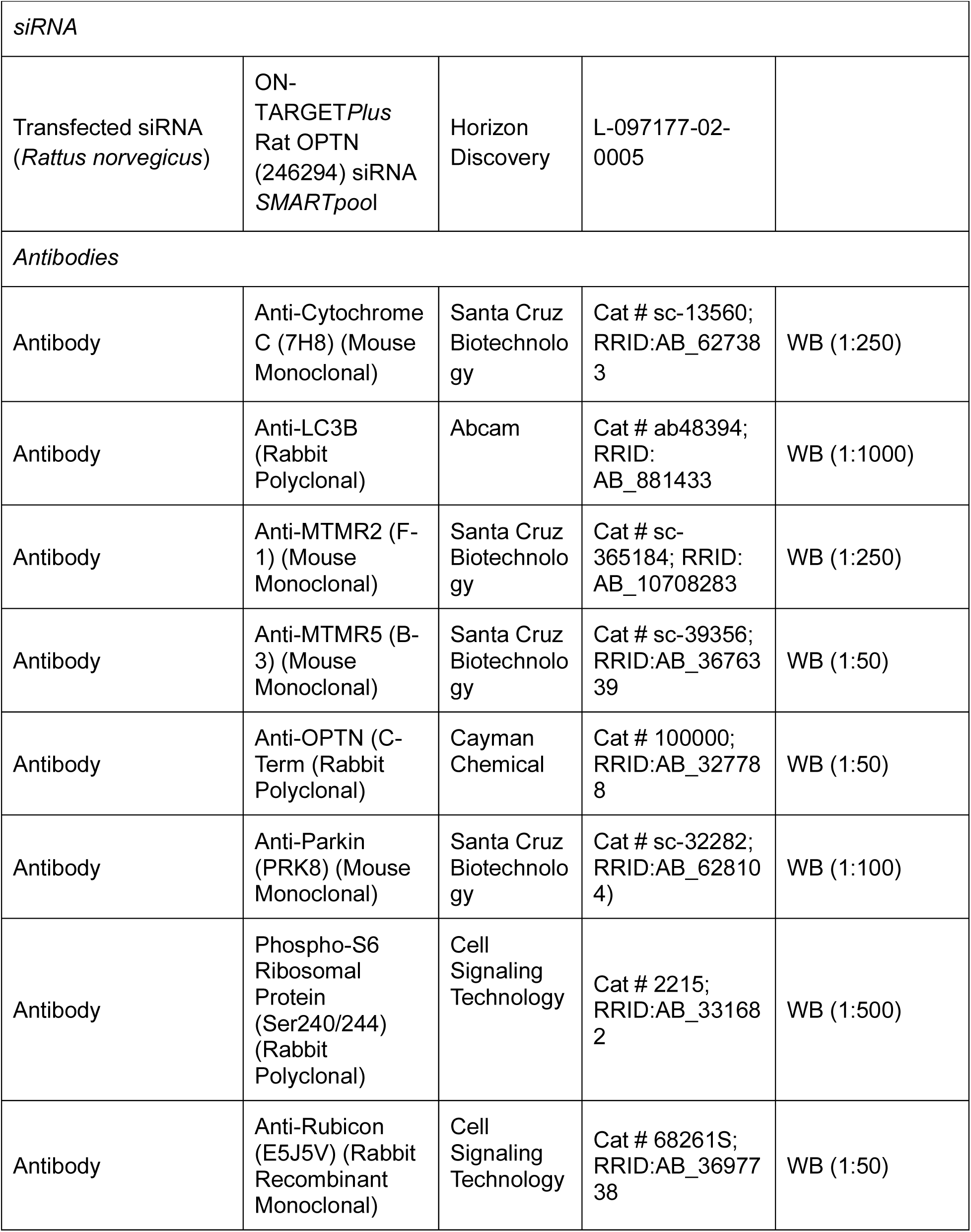
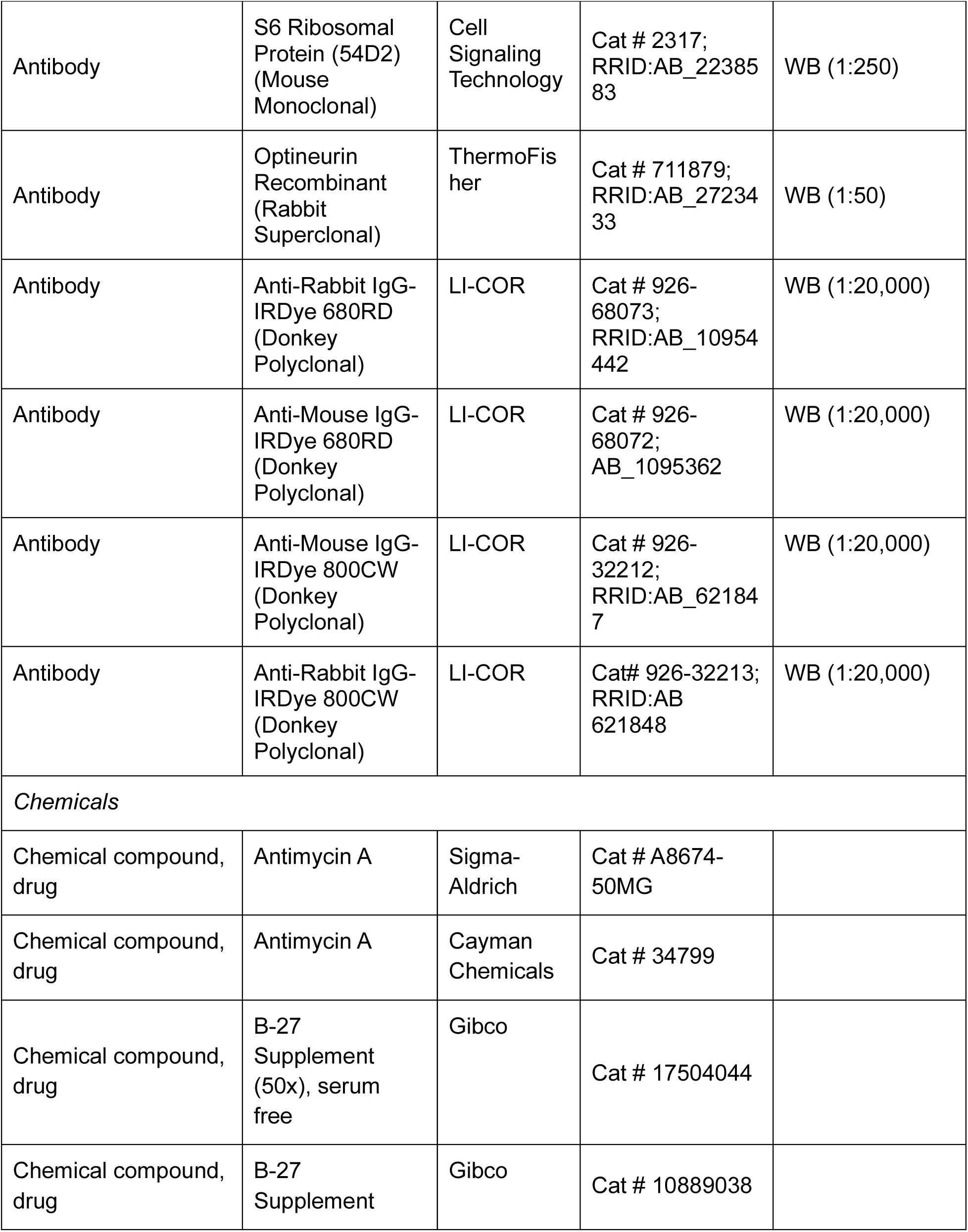
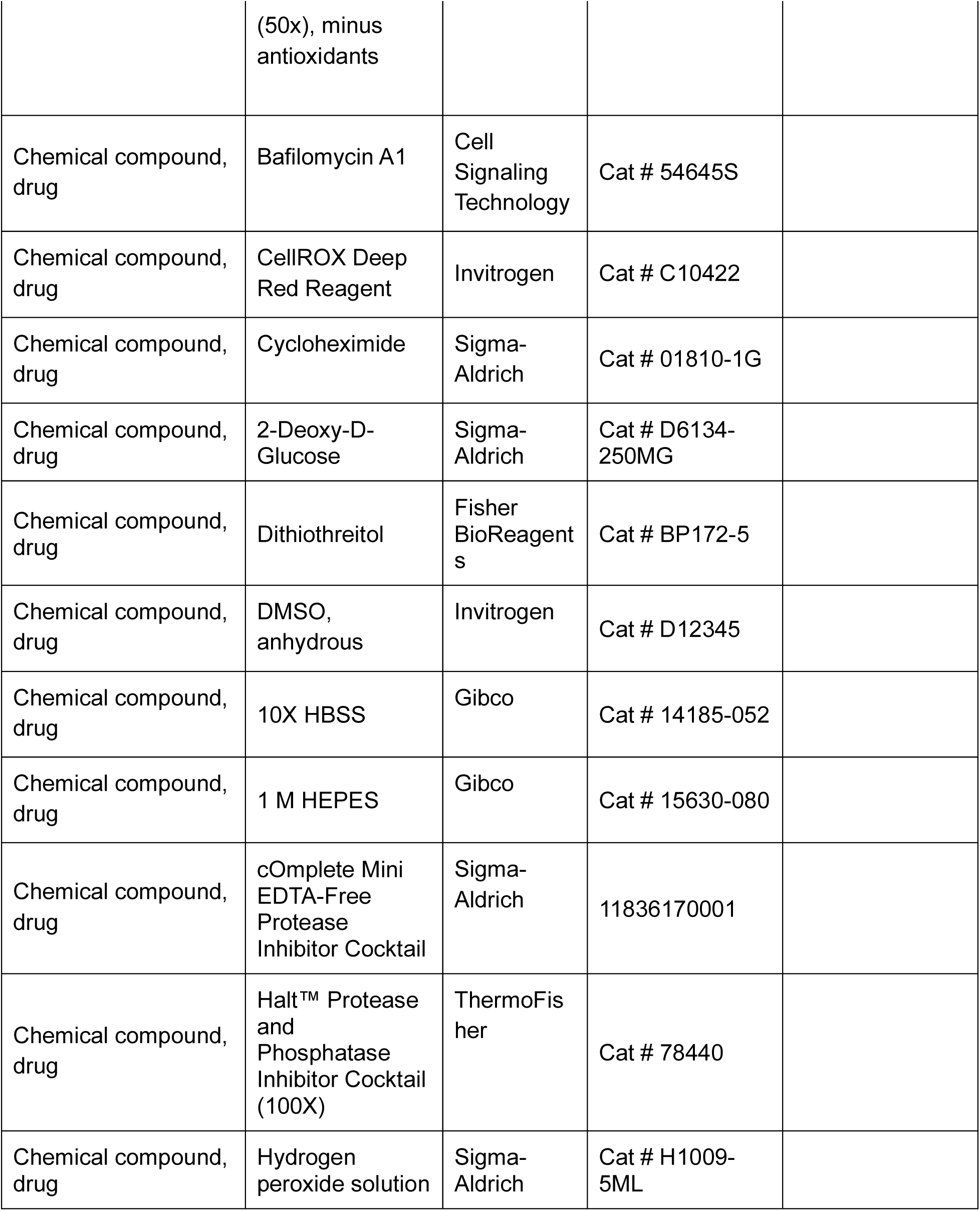
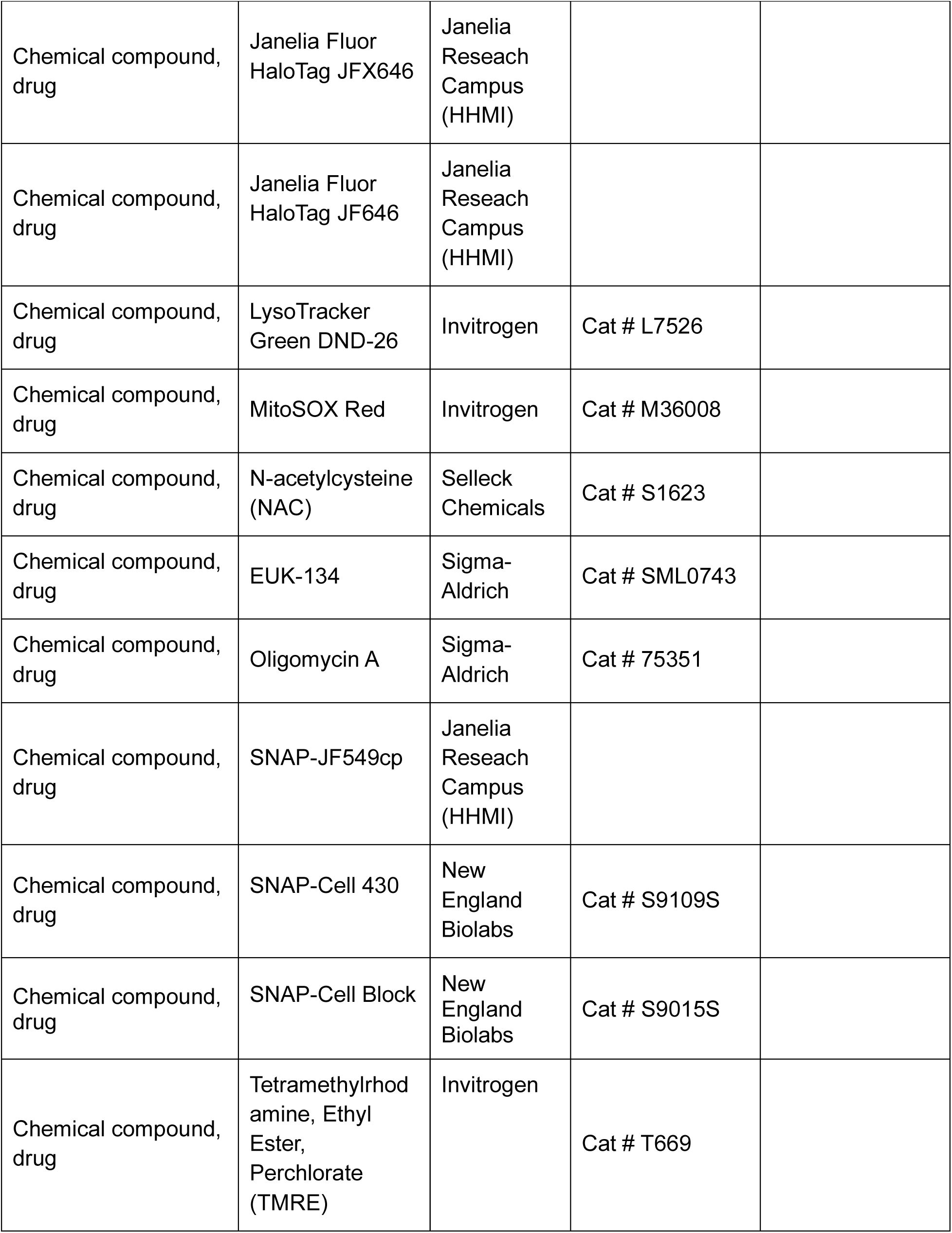
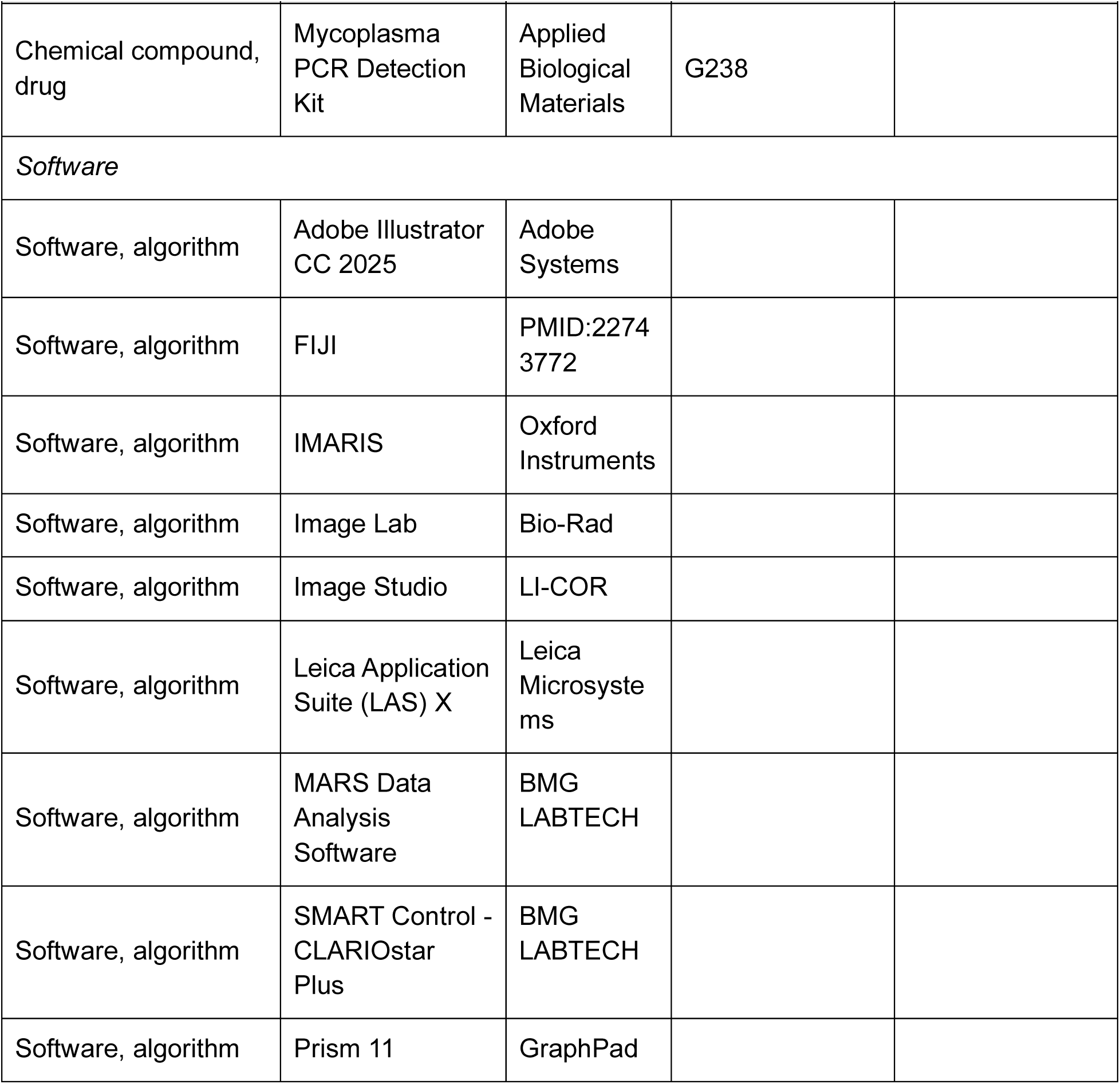
Key Resources Table.

**Figure S1:**
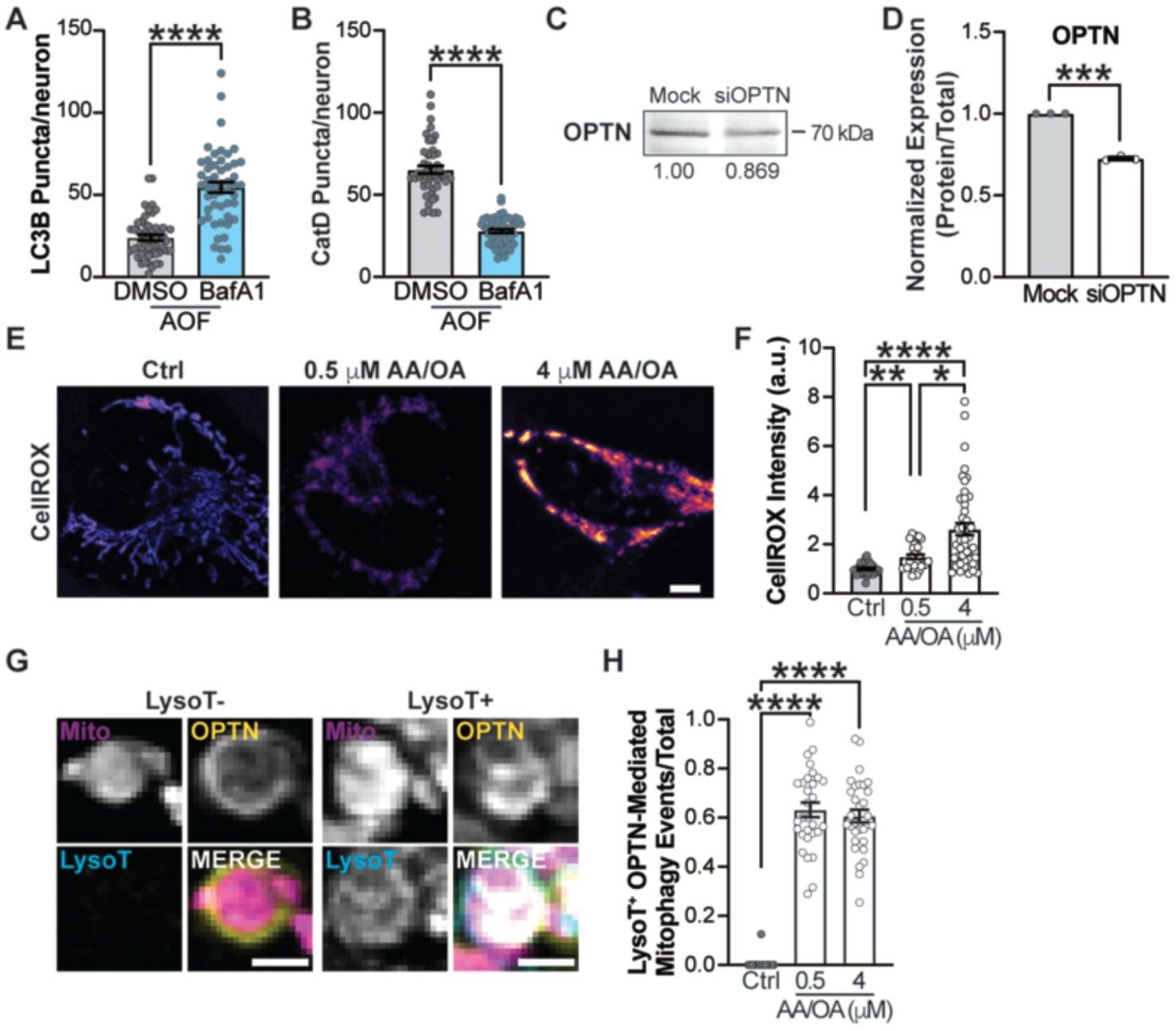
The levels of ROS do not alter the fast rates of mitochondrial turnover in HeLa cells. (**A**) Quantification of the number of LC3B+ puncta per neuron in the soma of hippocampal neurons following a 6-hour AOF treatment in either DMSO or BafA1. Mean ± SEM; *n* = 50-51 neurons from 5 biological replicates; 8-9 DIV. ****, *p* < 0.0001 by unpaired Mann-Whitney test. (**B**) Quantification of CatD+ puncta per neuron in the soma of hippocampal neurons following a 6-hour AOF treatment in either DMSO or BafA1. Mean ± SEM; *n* = 49-52 from 5 biological replicates; 8-9 DIV. ****, *p* < 0.0001 by unpaired Mann-Whitney test. (**C-D**) Representative fluorescent western blot (C) and quantification (D) of OPTN in neuron lysates. Data shown as the fold change over control of the protein of interest divided by the total protein stain. Normalization factors are shown under the corresponding representative images. Mean ± SEM; *n* = 3 biological replicates; 9 DIV. ***, *p* < 0.001 by Welch’s test. (**E-F**) Representative images (E) and quantification (F) of intracellular ROS using CellROX Deep Red. HeLa cells were treated for 2 hours with either control, 0.5 µM AA/0.5 µM OA, or 4 µM AA/4 µM OA. Scale bar, 5 µm. Mean ± SEM; *n* = 33-46 cells from 3 biological replicates. *, *p* < 0.05; **, *p* < 0.01; ****, *p* < 0.0001 by Kruskal-Wallis with Dunn’s multiple comparisons test. (**G-H**) Representative images (G) and quantification (H) of the fraction of OPTN-mediated mitophagy events that are LysoT+ following a 2-hour treatment. Scale bar, 1 µm. Mean ± SEM; *n* = 30-32 cells from 3 biological replicates. ****, *p* < 0.0001 by Kruskal-Wallis with Dunn’s multiple comparisons test.

**Figure S2:**
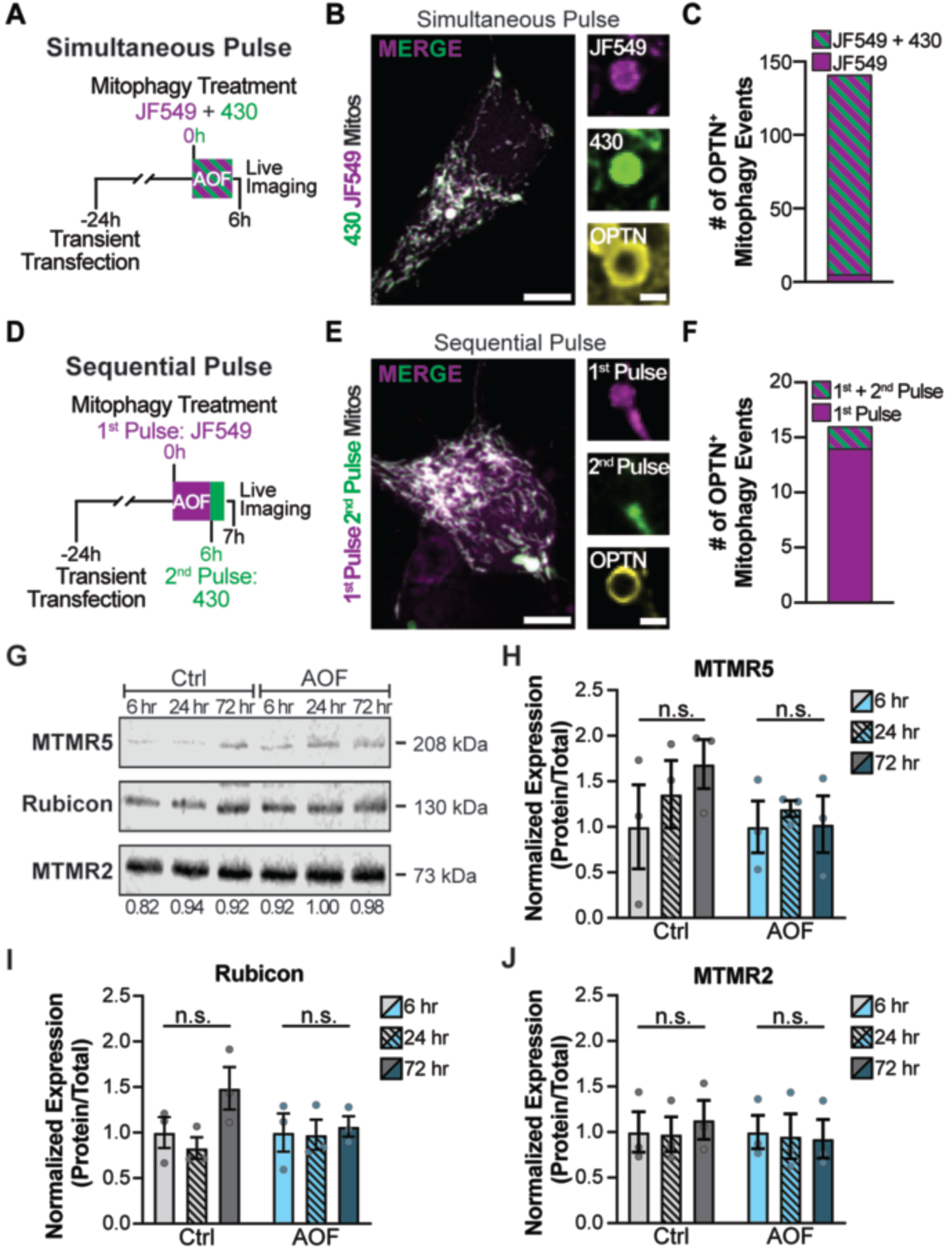
Negative autophagy regulators’ abundance does not change throughout the pulse-chase paradigm. (**A**) Experimental timeline for simultaneous SNAP pulse-chase labeling with SNAP JF549 and SNAP 430. Briefly, transiently transfected neurons were treated with AOF for 6 hours to induce OPTN-dependent mitophagy and simultaneously labeled with two SNAP ligands. After the treatment ended, the neurons were immediately imaged. (**B**) Representative images of the somal mitochondrial network (left) and OPTN+ mitophagy event (right) following simultaneous SNAP labeling. Scale bar: 5 µm (left); 1 µm (right). (**C**) Quantification of the total number of OPTN+ mitophagy events that are single-labeled (JF549) or dual-labeled (SNAP JF549 + 430). *n* = 25 neurons from 3 biological replicates; 7 DIV. (**D**) Experimental timeline for sequential SNAP pulse-chase labeling. Briefly, transiently transfected neurons were treated in parallel with AOF for 6 hours and labeled with SNAP JF549. Next, neurons were immediately labeled with SNAP 430 for 1 hour and imaged. (**E**) Representative images of the mitochondrial network (left) and OPTN+ mitophagy event (right) following sequential SNAP pulses. Scale bar: 5 µm (left); 1 µm (right). (**F**) Quantification of the total number of OPTN+ mitophagy events labeled with the 1^st^ Pulse (SNAP JF549) or 1^st^ + 2^nd^ Pulse (dual-labeled with SNAP JF549 and SNAP 430) when ligands were added sequentially. *n* = 31 neurons from 3 biological replicates; 7 DIV. (**G-J**) Representative fluorescent western blot (G) and quantification of MTMR5 (H), Rubicon (I), and MTMR2 (J) following 6-, 24-, or 72-hour recovery after control or AOF treatment. Data are shown as the fold change over the mean of the 6-hour recovery protein of interest divided by the total protein stain. Normalization factors are shown under the corresponding representative images. Mean ± SEM; *n* = 3 biological replicates; 7 DIV. n.s., not significant by One-way ANOVA with Tukey multiple comparisons test (H-I) and Brown-Forsythe and Welch’s ANOVA with Dunnett T3 multiple comparisons (J).

**Figure S3.**
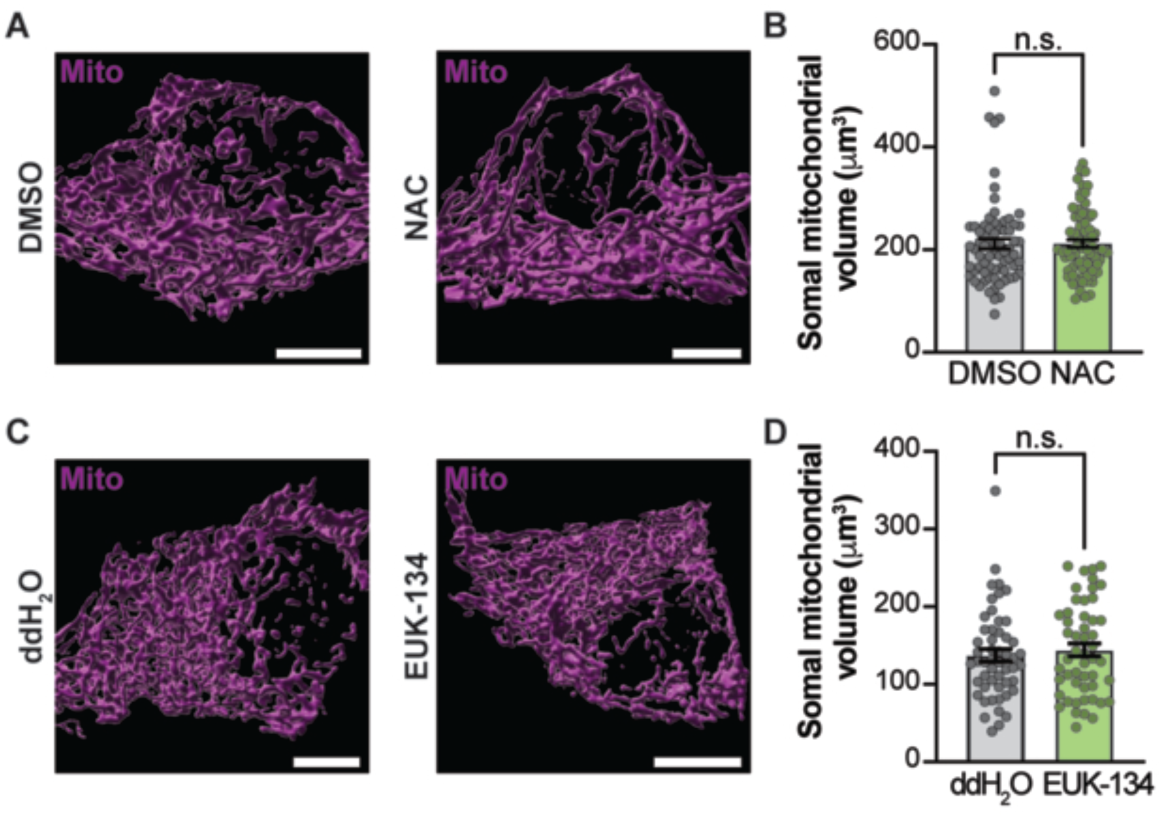
Treatment with NAC or EUK-134 does not alter mitochondrial volume. (**A-B**) Representative images (A) and quantification (B) of somal mitochondrial surface volumes in DMSO-treated (left) and NAC-treated (right) neurons. Scale bar, 5 µm. Mean ± SEM; *n* = 70 neurons from 4 biological replicates; 8 DIV. n.s., not significant by unpaired Mann-Whitney test. (**C-D**) Representative images (C) and quantification (D) of somal mitochondrial surface volumes in ddH_2_O-treated (left) and EUK-134-treated (right) neurons. Scale bar, 5 µm. Mean ± SEM; *n* = 51-52 neurons from 3 biological replicates; 8 DIV. n.s., not significant by unpaired Mann-Whitney test.

**Figure S4:**
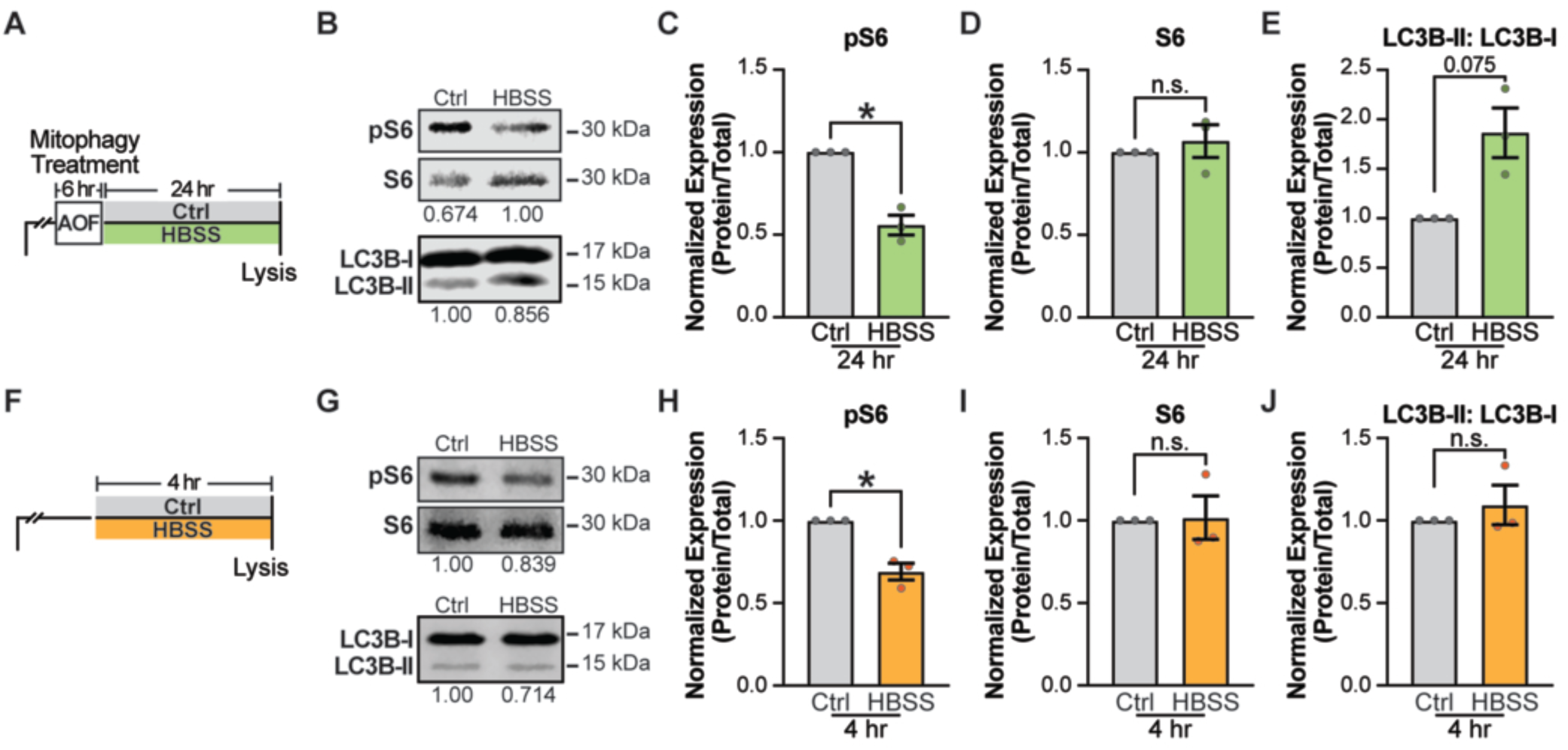
LC3B lipidation is enhanced after 24 hours, but not 4 hours, of partial nutrient deprivation. (**A**) Schematic illustrating the neuronal starvation experimental timeline. Briefly, a 6-hour AOF treatment induces OPTN-mediated mitophagy, followed by 24 hours in either control or HBSS to induce partial nutrient deprivation. (**B-E**) Representative fluorescent western blot (B) and quantification of pS6 (C), S6 (D), and LC3B-II:LC3B-I ratio (E) in neuron lysates. Data shown as the fold change over control of the protein of interest divided by the total protein stain. Normalization factors are shown under the corresponding representative images. Mean ± SEM; *n* = 3 biological replicates; 8-9 DIV. n.s., not significant; *, *p* < 0.05; *p* = .0747 by Welch’s t-tests. (**F**) Schematic of the short partial nutrient deprivation timeline, where neurons were treated for 4 hours with either control or HBSS. (**G-J**) Representative fluorescent western blot (G) and quantification of pS6 (H), S6 (I), and LC3B-II:LC3B-I ratio (J) in neuron lysates. Data shown as the fold change over control of the protein of interest divided by the total protein stain. Normalization factors are shown under the corresponding representative images. Mean ± SEM; *n* = 3 biological replicates; 8 DIV. n.s., not significant; *, *p* < 0.05 by Welch’s t-tests.

**Figure S5:**
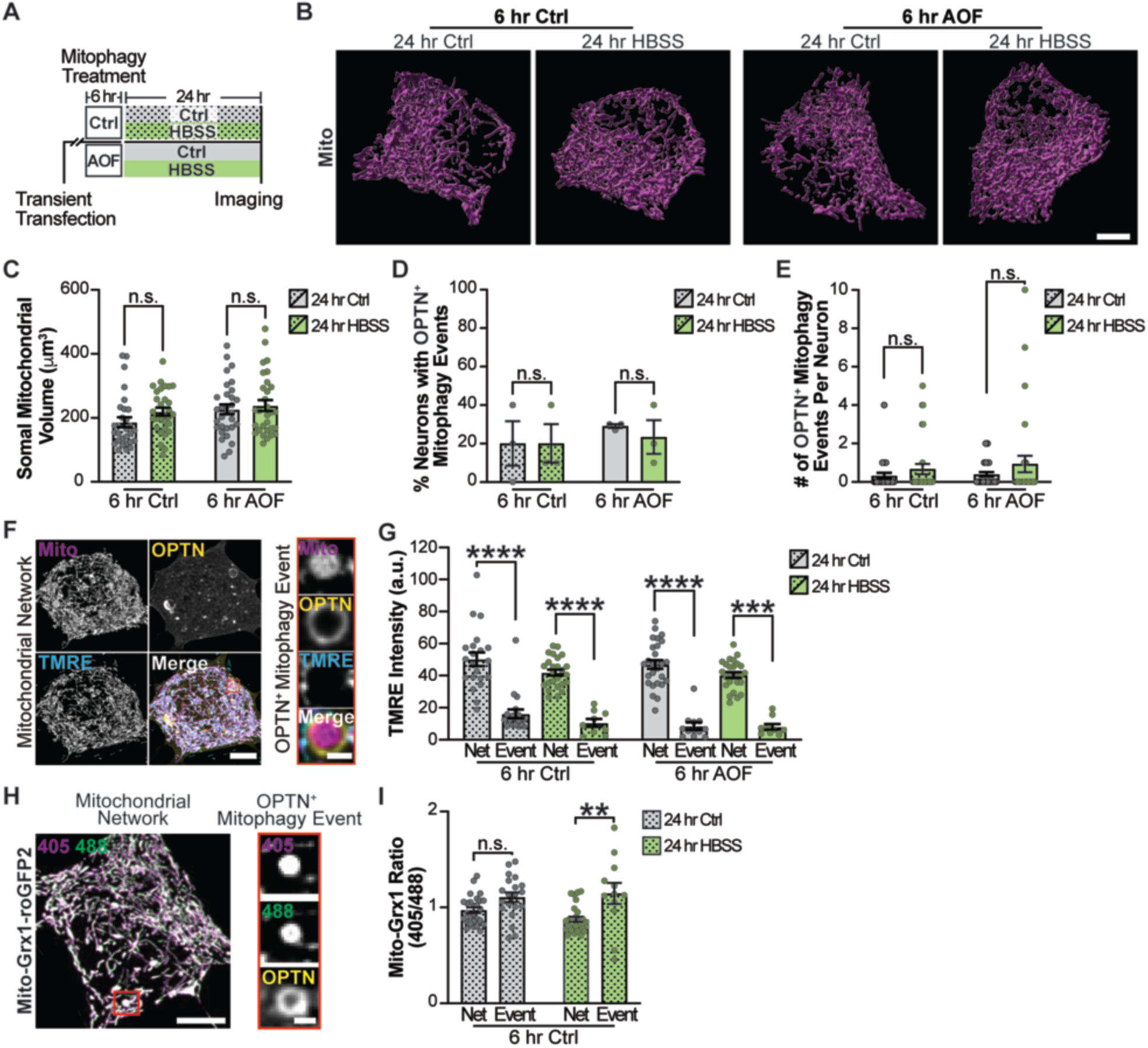
Partial nutrient deprivation does not affect the mitochondrial network or mitophagy induction. (**A**) Schematic illustrating a longer partial nutrient deprivation timeline. Neurons were treated for 6 hours with control or AOF, followed by 24 hours in either control or HBSS. (**B-C**) Representative images (B) and quantification (C) of somal mitochondrial surface volumes. Mean ± SEM; *n* = 29-31 neurons from 3 biological replicates; 8-9 DIV. n.s., not significant by Two-Way ANOVA with Šídák’s multiple comparisons test. (**D**) Quantification of the percent of neurons undergoing OPTN-mediated mitophagy. Mean ± SEM; *n* = 9-28 events in 29-31 neurons from 3 biological replicates; 8-9 DIV. n.s., not significant by Two-Way ANOVA with Šídák’s multiple comparisons test. (**E**) Quantification of the number of OPTN+ mitophagy events per neuron. Mean ± SEM; *n* = 9-28 events in 29-31 neurons from 3 biological replicates; 8-9 DIV. n.s., not significant by Two-Way ANOVA with Šídák’s multiple comparisons test. (**F-G**) Representative images (F) and quantification (G) of TMRE in the mitochondrial network and at OPTN+ mitophagy events. Mean ± SEM; *n* = 9-20 events in 21-27 neurons from 3 biological replicates; 9-10 DIV. *** *p* < 0.001; **** *p* < 0.0001 by Kruskal-Wallis test with Dunn’s multiple comparison test. (**H-I**) Representative images (H) and quantification (I) of Mito-Grx1-roGFP2 within the mitochondrial network and at OPTN+ mitophagy events. Scale bar, 5 µm (Left); 1 µm (Right). Mean ± SEM; *n* = 13-20 events from 3 plates per condition, which includes 27-28 neurons across 3 biological replicates; 9 DIV. n.s., not significant; **, *p* < 0.01 by Two-Way ANOVA with Tukey’s multiple comparisons test. Net, network.

